# Using XCiMS to Quantify the Benefits of Ion Mobility Spectrometry for Untargeted Metabolomics

**DOI:** 10.64898/2026.10.21.760801

**Authors:** Lingjue Wang, Kevin Cho, Junjie Yang, Ethan Stancliffe, Michaela Schwaiger-Haber, Clark S. Hofstrand, Triston Groff, James N. Dodds, Leah P. Shriver, Dhanalakshmi S. Anbukumar, Erin S. Baker, Gary J. Patti

## Abstract

Integrating ion mobility spectrometry (IMS) with liquid chromatography-mass spectrometry (LC-MS) has the potential to improve untargeted metabolomics workflows, but assessing the benefits of IMS has been challenging due to a paucity of software solutions to process raw LC-IMS-MS data in a manner that enables direct comparison with LC-MS data. We developed the XCiMS software to collapse 4D data into pseudo-3D data. Peak detection is then performed in the pseudo-3D space, followed by targeted extraction of 4D signals from 3D-guided regions of interest. Here, we applied XCiMS to estimate the potential of IMS for untargeted metabolomics when analyzing human plasma. First, we assessed the number of metabolites that are uniquely resolved by LC-IMS-MS but not LC-MS. Although as many as 50% of 3D peaks produce multiple 4D peaks when using high-resolution IMS, few represent newly resolved structural isomers. We found that the increase in features due to IMS was mostly caused by charge isomers and ion clusters. Second, we investigated the impact of IMS on metabolite identification. When using a 16-minute chromatographic separation, we found that IMS can decrease the frequency of chimeric MS/MS by about half. We conclude that while IMS adds considerable complexity to untargeted metabolomics data without resolving a large number of additional structural isomers relative to a standard LC-MS experiment, it does facilitate metabolite identification by decreasing chimeric spectra and providing CCS values for library searching.

## Introduction

The objective of untargeted metabolomics is to profile as many small molecules in a biological system as possible.^1^ At present, liquid chromatography-mass spectrometry (LC-MS) is widely regarded as the leading analytical platform best suited to achieve this goal.^2, 3^ Nevertheless, even LC-MS falls short of achieving comprehensive coverage of the metabolites, lipids, and chemical exposures present in a biological sample.^4, 5^ Consequently, the search for improved technologies capable of detecting a broader range of small molecules remains a major research focus in metabolomics.^6, 7^

Ion mobility spectrometry (IMS) offers a promising strategy to extend the chemical space accessible by LC-MS.^8–13^ IMS differentiates ions in the gas phase on the basis of their size, shape, and charge at a timescale that is compatible with both chromatographic separation and rapid mass spectrometric acquisition (**Figure 1a**).^14^ Given the large number of isomeric small molecules, the integration of IMS with LC-MS provides an opportunity to enhance untargeted metabolomics by adding an orthogonal dimension of molecular separation.^12^ Over the past two decades, LC-IMS-MS platforms have become increasingly accessible with the introduction of commercial instruments from Waters, Agilent, and Bruker. Despite the availability of the technology, an important question in the field remains: to what extent does the incorporation of IMS into LC-MS workflows improve untargeted metabolomics?

**Figure 1.**
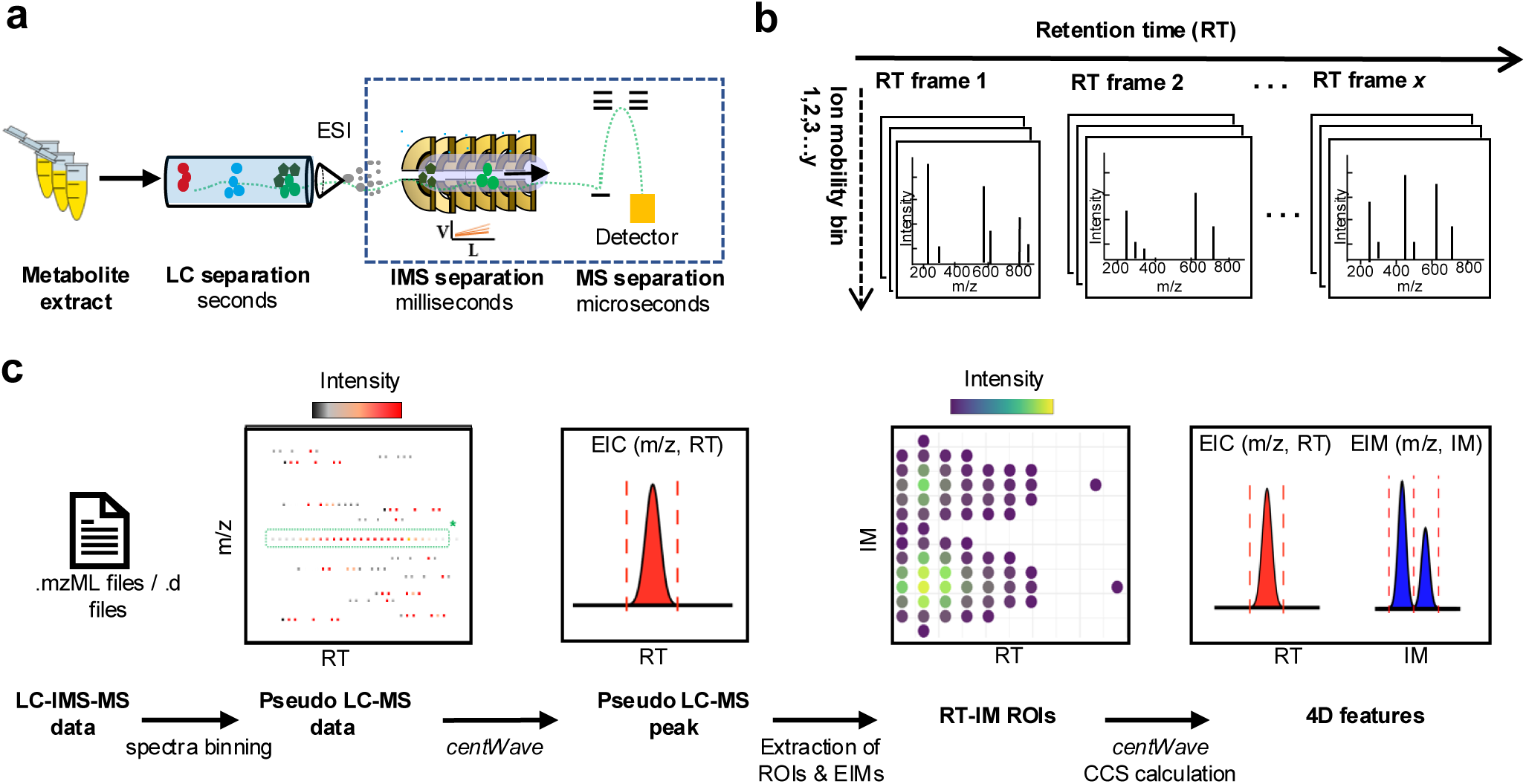
Overview of LC-IMS-MS analysis and XCiMS workflow. a) General schematic of LC-IMS-MS. b) Data acquisition occurs across retention time frames and ion mobility bins. c) The first step of the XCiMS workflow is to bin mass spectra from LC-IMS-MS data files to create pseudo LC-MS data for peak detection, resulting in extracted ion chromatograms (EIC, red). At regions of interest (ROIs), IM data are extracted to create extracted ion mobilograms (EIM, blue).

To date, it has been challenging to quantify the benefits of IMS in untargeted metabolomics because of limitations in computational approaches for processing and interpreting 4D data. While important progress has been made^15–18^, far fewer informatics tools are available for LC-IMS-MS data than for conventional LC-MS analyses. Widely used software platforms, such as XCMS^19, 20^, have enabled 3D workflows but were not designed to accommodate the additional dimension provided by IMS. Here, we introduce XCiMS to address this gap. The unique processing pipeline of XCiMS collapses LC-IMS-MS data into pseudo-3D LC-MS data. Beyond enabling robust quantitative profiling, our approach allows users to compare 3D and 4D metabolomics data directly. The application of XCiMS to human plasma analyzed by three different IMS technologies revealed that 4D profiling adds considerable complexity to the data without resolving a large number of additional structural isomers compared to conventional LC-MS workflows. On the other hand, our analysis showed that IMS facilitates metabolite identification by reducing the number of chimeric MS/MS spectra approximately 50%. Chimeric MS/MS spectra are defined as MS/MS spectra containing signals derived from two or more precursor ions that are isolated and fragmented simultaneously. XCiMS is freely available as an R package via GitHub: pattilab.github.com/XCiMS.

## Results

### Developing the XCiMS workflow

We developed XCiMS to process LC-IMS-MS data, which we refer to throughout this work as 4D data, within the XCMS ecosystem. We validated that the software is compatible with data acquired from Agilent DTIMS, Waters TWIMS, and Bruker TIMS instruments. The XCiMS workflow first constructs vendor-neutral LC-IMS-MS spectra by parsing centroided raw data from .mzML or TIMS .d files. The 4D spectra are partitioned into two orthogonal dimensions, denoted as retention time (RT) frames and ion mobility (IM) bins (**Figure 1b and Supplementary Figure 1a)**. After collapsing the data into a pseudo-3D format (**Figure 1c and Supplementary Figure 1b**), XCiMS applies the *centWave* algorithm to detect 3D precursors.^21^ For each detected 3D feature, an extracted ion mobilogram (EIM) is then constructed by summing all MS peaks for each IM bin that fall within a user-specified *m/z* tolerance of the detected 3D feature and across the retention time where the 3D feature elutes (**Supplementary Figure 1c-d**). This operation retains the peak profiles and improves the signal to noise. Lastly, peak detection of the EIMs is performed by using the *centWave* algorithm. The resulting peaks are merged with their corresponding 3D precursors to generate a table of 4D features. XCiMS has the capability to process multiple data files, perform correspondence determination, and calculate collision cross-section (CCS) values for various types of IMS technologies. A detailed description of each function in the XCiMS workflow is included in the **Methods**.

We consider a 4D feature to be an ion signal with a unique combination of *m/z*, retention-time, and ion mobility (IM) values. Inspired by the terminology used in mathematics^22^, we define two types of 4D features: injective and non-injective 4D features (**Supplementary Figure 2**). We use injective to refer to 4D features that map directly back to 3D features in a one-to-one fashion (i.e., one 3D feature produces one 4D feature). In contrast, when multiple 4D features originate from a single 3D feature, the one-to-one correspondence is lost and we classify all of the associated 4D features as non-injective. The injective and non-injective nomenclature is convenient for recognizing when IMS may be resolving structural isomers. For example, structural isomers that co-elute in the LC dimension but separate in the IMS dimension will yield non-injective 4D features.

### Quantitative Performance of XCiMS

To evaluate the quantitative performance of XCiMS for 4D metabolomics, we analyzed 13 authentic amino acids (AAMix) by hydrophilic interaction liquid chromatography (HILIC)-TWIMS-MS in negative mode. The dataset was first manually curated by using Skyline.^23^ We confirmed that [M-H]^-^ions of the 13 amino acids were detected across three technical replicates, yielding a total of 39 reference signals. To process the data in an automated fashion by using XCiMS, we optimized *centWave* parameters so that the maximum number of amino acid reference signals were detected. Using the adjusted parameters, XCiMS detected all expected signals. Examples of extracted ion chromatograms and ion mobilograms from XCiMS and Skyline are shown in **Figure 2a**. All 13 amino acids detected by XCiMS are shown in **Supplementary Figure 3**.

**Figure 2.**
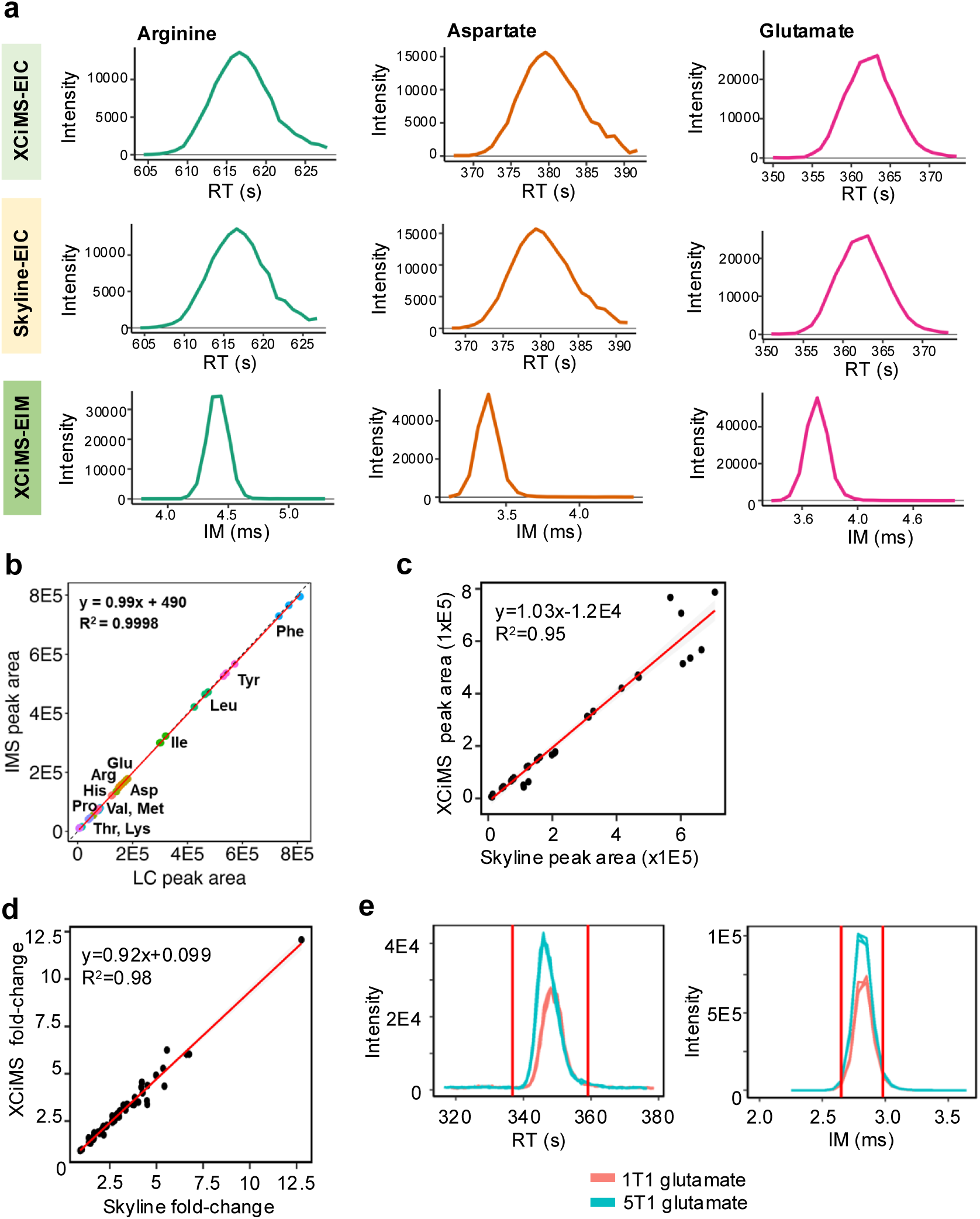
Quantitative performance of XCiMS. a) Comparison of peak fidelity between XCiMS and Skyline. Representative extracted ion chromatograms (EICs) and extracted ion mobilograms (EIMs) from XCiMS and Skyline. b) Correlation between LC and IMS peak areas of 13 amino acid standards as detected by XCiMS. c) correlation of peak areas obtained from XCiMS and Skyline. d) XCiMS-detected fold changes are highly correlated with manually calculated fold changes from Skyline-integrated peak areas. Data from 57 4D features with a p value <0.05 between the 1:1 and 5:1 spiked samples are shown. e) Pseudo LC-MS peak and IMS peak for glutamate as a representative example of data from the spike-in experiment. Note that human plasma already contains glutamate. Accordingly, the difference in glutamate intensity between the spike-in samples is not expected to be 5.

In principle, the peak area of an injective 4D feature and the peak area of its corresponding 3D feature should be equal. For non-injective 4D features, the area of a 3D peak should equal the sum area of all of the 4D peaks derived from it. In this particular analysis of amino acid standards, all 4D features were injective (i.e., only one 4D peak was detected for each amino acid). As such, we expected IM and LC peak areas to be comparable. Indeed, XCiMS returned IM and LC peak areas that were identical (**Figure 2b**). We also compared the peak areas returned by XCiMS to those that were manually integrated in Skyline. The results show a strong correlation, suggesting accurate and reproducible multidimensional peak detection by XCiMS (**Figure 2c**).

To demonstrate the ability of XCiMS to measure fold changes between samples, a spike-in experiment was performed. Two groups of samples were prepared by spiking a mixture of polar metabolites (PolarMix) into SRM1950 human plasma at different concentrations. The first group of samples was made by spiking 1 equivalent of the PolarMix into SRM1950 plasma extract (labeled "1T1"), while the second group of samples was made by spiking 5 equivalents of the PolarMix into the same volume of SRM150 plasma extract (labeled "5T1"). The samples were analyzed by HILIC-TWIMS-MS in negative mode. Skyline, which allows for targeted processing of 4D data, was used to obtain peak areas for the reference [M-H]^-^signals of the 65 standards. Automated peak detection, feature correspondence, and fold-change analysis was accomplished in parallel by using XCiMS (**Methods**). XCiMS successfully detected signals for each of the 65 standards spiked in the samples. Given that the standards in the spike-in mix were also endogenously present in human plasma, the fold change of each compound varied between groups. Notwithstanding, the fold changes output by XCiMS were consistent with the fold changes calculated from peak areas in Skyline, demonstrating that XCiMS accurately assesses fold changes between sample groups (**Figure 2d**). Glutamate as an example of a spiked metabolite is shown in **Figure 2e**.

### Using XCiMS to Optimize LC-IMS-MS Workflows

When designing a targeted analysis for a single compound, it is relatively straightforward to optimize a method.^24^ Most often, a chemical standard is evaluated over a range of experimental conditions to test which provides optimal results. For untargeted metabolomics studies, on the other hand, the goal is to profile the entire collection of small molecules within a biological system. In such cases, it is impractical to obtain chemical standards for every compound of interest. Historically, one alternative strategy has been to identify protocols that maximize the number of features detected by programs such as XCMS.^25^ In this sense, data-processing platforms have played a key role in optimizing experimental conditions for untargeted metabolomics, ranging from sample-handling protocols to mass spectrometry settings.^26–30^

A barrier to advancing LC-IMS-MS for untargeted metabolomics has been the paucity of software solutions to evaluate data at a comprehensive scale. We expect XCiMS to help close this gap. As an example, we applied XCiMS to optimize instrument settings when using LC-TWIMS-MS. Rather than using total features as a metric of performance, we applied a strategy known as credentialing.^25^ As we have demonstrated previously, credentialing introduces isotopic signatures into the data that enable non-biological signals to be filtered, thereby providing a more accurate estimate of metabolite coverage.^31^ Credentialed *E. coli* samples were prepared as described before^32^ and detailed in Methods. In brief, *E. coli* samples were cultured in ^12^C or ^13^C media and then mixed at 1:1 and 1:2 ratios prior to being analyzed by LC-IMS-MS. For a feature to be credentialed, it must appear in mass spectra as a doublet having ^12^C and ^13^C species at the appropriate ratio.

We considered three TWIMS-MS settings: wave height, wave velocity, and resolution mode. Experiments were performed by using three distinct sets of parameters (**Supplementary Figure 4a-c**). The first set of parameters returned 2815 credentialed 3D features and the second set of parameters returned 2844 credentialed 3D features. Interestingly, the number of credentialed 4D features increased from 1917 with parameter set 1 to 3255 with parameter set 2, indicating a substantial improvement in TWIMS resolution (**Supplementary Figure 4b**). An even larger increase in features was observed when we changed the resolution mode. Contrary to intuition, we observed a 470% and 660% increase in 3D and 4D features, respectively, when we used standard resolution mode instead of high-resolution mode (**Supplementary Figure 4c)**. Our interpretation of these findings is that a mismatch between high ion traveling speeds in TWIMS and slower data sampling rates by the TOF in high-resolution mode negatively affects data quality in the IMS dimension. Nonetheless, the results demonstrate that XCiMS can be leveraged to optimize LC-IMS-MS workflows for discovery analysis.

### Sources of non-injective features reported by XCiMS

As noted earlier, LC-IMS-MS can generate two types of 4D features: injective and non-injective (**Supplementary Figure 2**). We considered three potential sources of non-injective 4D features in our data: (i) structural isomers, (ii) charge isomers, and (iii) dissociation of ion clusters post-IMS separation. Structural isomers lead to non-injective 4D features when they co-elute chromatographically but are then resolved by IMS. This possibility presents an exciting opportunity for untargeted metabolomics, where IMS differentiates chemical species that potentially yield important biological insight. A second source of non-injective 4D features is charge isomers, which are ions that have the same elemental composition but differ in the physical location of the charge within the molecule.^33^ The site of the charge can influence the shape and conformation of a molecule, thereby giving it a unique CCS value.^34^ A third source of non-injective 4D features is the dissociation of ion clusters. As we and others have reported previously, co-eluting metabolites can form complex ion clusters at the source or along the ion path.^35, 36^ When these ion clusters dissociate after IMS separation, they produce signals with the same *m/z* values as their individual monomer units but with apparently different CCS values.

To demonstrate that non-injective 4D features reported by XCiMS can originate from any of the three sources outlined above, we sought to identify examples of each. The examples described in this section were identified manually by using reference data from authentic chemical standards. Using our HILIC method, we found that the structural isomers glucose 1-phosphate (G1P) and fructose 6-phosphate (F6P) can only be differentiated by IMS (**Figure 3a**). While G1P and F6P are structurally similar, they are involved in distinct biochemical processes. G1P is associated with glycogen metabolism, whereas F6P is an intermediate of glycolysis.^37^ Their separation by IMS, but not LC, highlights a benefit of profiling metabolites in 4D compared to 3D. The non-injective features associated with adenosine, on the other hand, showcase the complexity that IMS adds to untargeted metabolomics data (**Figure 3b**). In positive mode, adenosine can be protonated at multiple sites within the molecule. At least two of those charge isomers are measured from our samples, as supported by reference data from the Unified CCS Compendium (**Supplementary Figure 5**).^38^ Similarly contributing to the complexity of 4D data is the dissociation of ion clusters, such as the heterodimer formed between cytidine and tryptophan (**Figure 3c**). The co-elution of cytidine and tryptophan results in the formation of an ion cluster, but the cluster dissociates after IMS separation. As a result, we measure the same *m/z* values for cytidine and tryptophan with different CCS values (**Supplementary Figure 6**).

**Figure 3.**
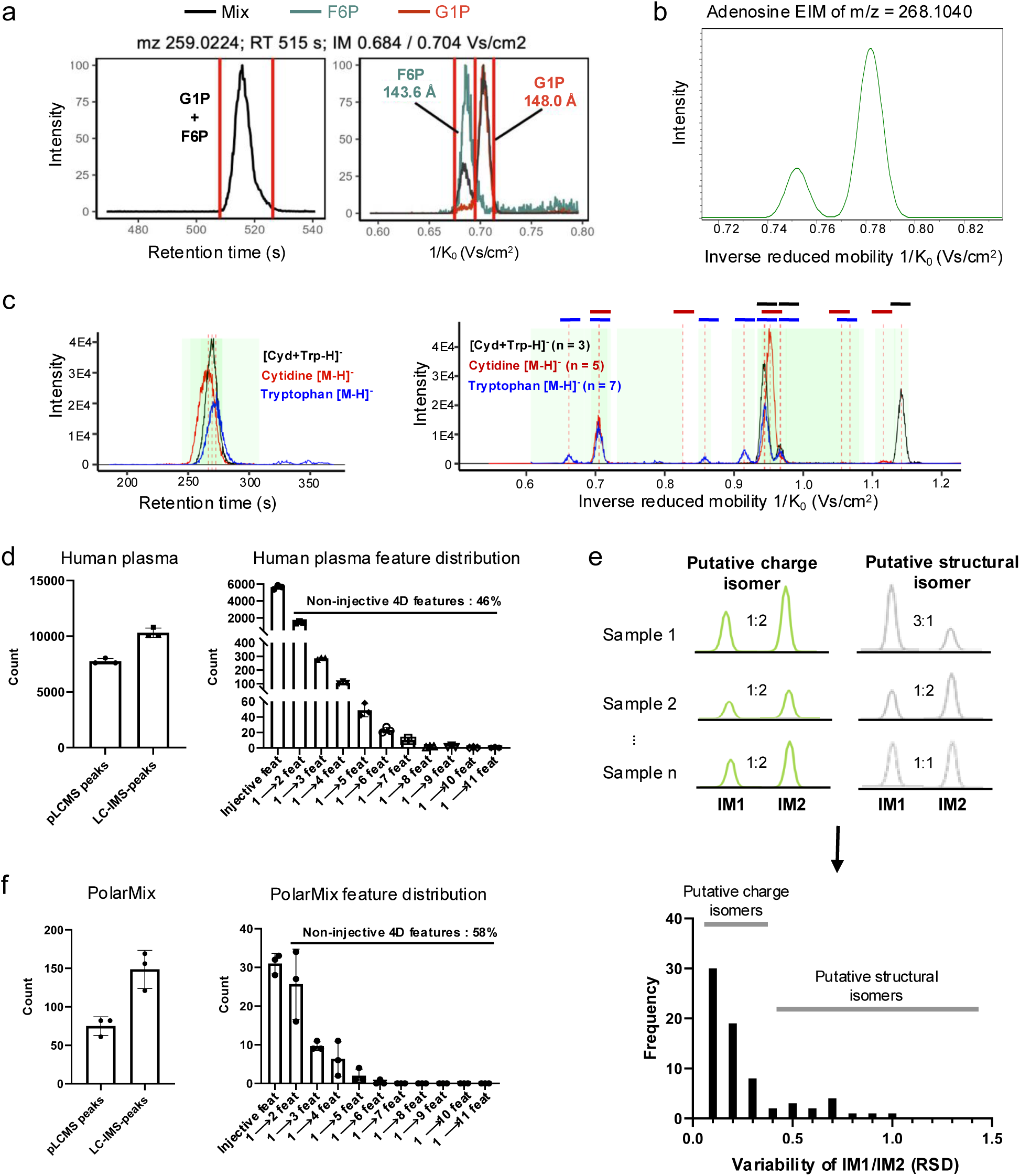
Non-injective 4D features. a) EIC for *m/z* 259.0224 shows co-elution of two hexose phosphates in the LC dimension. IMS separates the two structural isomers, resolving fructose 6-phosphate (F6P) and glucose 1-phosphate (G1P). The black line represents signal from the PolarMix. The green and red lines represent signals from the analysis of pure F6P or G1P standards, respectively. Each plot is normalized to 100. b) Two IMS peaks were detected for adenosine from research samples and an authentic chemical standard for the [M+H]+ ion. More details are shown in **Supplemental Figure 5**. c) Example of an ion cluster leading to multiple 4D peaks. Pseudo LC-MS peaks and IMS peaks as reported by XCiMS for cytidine [M-H]-(red), tryptophan [M-H]-(blue), and a cytidine-tryptophan ion cluster [Cyd+Trp-H]-(black). The green rectangles and red dashed vertical lines indicate boundaries and apexes of the chromatographic and IMS peaks detected by XCiMS. More details are shown in **Supplemental Figure 6.** d) Number of pseudo LC-MS (pLC-MS) peaks (3D) and LC-IMS-MS peaks (4D) detected from human plasma extract by using LC-TIMS-MS. The distribution of 4D features shows a large number of non-injective features. e) Strategy to distinguish potential charge isomers from structural isomers by using XCiMS. Different biological samples are measured. When the variability of the ratio between two noninjective 4D features is low across different samples, the two features are more likely charge isomers. f) Number of pseudo LC-MS peaks and LC-IMS-MS peaks detected from a polar metabolite standard mix arising from chemical standards by using LC-TIMS-MS. Distribution of 4D features shows that a large number of non-injective features were also detected.

### Most non-injective features detected by XCiMS are not structural isomers

One motivation for incorporating IMS into the untargeted metabolomics workflow is to improve molecular resolution by differentiating structural isomers that cannot otherwise be distinguished by LC-MS alone. To make a quantitative assessment of the extent to which IMS accomplishes this goal, we need to determine the number of non-injective 4D features that originate from structural isomers. In theory, the evaluation could be conducted by identifying each of the features output by XCiMS with chemical structures. In practice, however, compound identification continues to be a bottleneck in untargeted metabolomics and assigning structures to each of the 4D features with high confidence is not practical. As an alternative, we devised experimental strategies to assess charge isomers and ion clusters without having to elucidate their chemical identities. Our logic is that, with an estimation of charge isomers and ion clusters in hand, we can then infer the remaining fraction of non-injective 4D features representing structural isomers.

Analysis of human plasma by LC-TIMS-MS yielded ∼8k 3D features and ∼11k 4D features (**Figure 3d**). The addition of TIMS resulted in a nearly 40% increase in feature count, meaning that almost half of the 3D features produced non-injective 4D features. Given that non-injective 4D features can arise from multiple sources, the results set an upper limit on the number of structural isomers that can be resolved by IMS but not LC with our 16-minute gradient. As a first step to improve our understanding of the data, we then analyzed human plasma by using a different IMS technology. Metabolites were extracted from human plasma with the same protocol and separated with identical chromatography, but instead of profiling by TIMS-MS, we used DTIMS-MS. Considering that the resolving power of TIMS and DTIMS were comparable in our experiments (∼50 based on singly charged ions, see Methods), we reasoned that co-eluting structural isomers would be similarly differentiated across platforms. Strikingly, while the number of 3D features measured by the different instruments was in the same ballpark, DTIMS resulted in far less non-injective 4D features. Less than 3% of 3D features produced more than one 4D feature with DTIMS, indicating that the number of structural isomers that can be resolved by an ion mobility system at a resolving power of 50 was less than a couple hundred.

Next, we aimed to better estimate the number of non-injective 4D features representing structural isomers in our DTIMS dataset. At the present time, there are no software tools to annotate charge isomers, which we suspected might be contributing to the non-injective 4D feature count. To assess charge isomers, we profiled plasma from eight human subjects and compared the ratios between co-eluting non-injective 4D features. The samples were processed and analyzed by using identical methods. We therefore assumed that the incidence at which charge isomers formed across samples would be similar, resulting in relatively constant ratios of co-eluting non-injective 4D features between different subjects. Structural isomers, on the other hand, are distinct metabolites whose levels are driven by biology. We surmised that the likelihood of two structural isomers having the same ratio across eight human subjects was low. As such, we assumed that structural isomers would produce non-injective 4D features whose ratios differed between human subjects (**Figure 3e**).

To validate our ratiometric approach, we considered some specific examples as test cases. Phenylalanine produced two 4D features in positive mode DTIMS analysis and provided data for a representative charge isomer. The ratio between the two 4D features was between 2.9 and 3.3 across the eight human subjects, with a variation of only 5.1%. The average technical variability of the same ratio from five replicate injections of a single sample was 9.8%. Using an authentic chemical standard, we confirmed that phenylalanine produced two 4D features at approximately a ratio of 3:1 (**Supplementary Figure 7a-b**). As a second test case for charge isomers, we used tryptophan (**Supplementary Figure 7c-d**). Tryptophan produced two 4D features that had a ratio between 1.6 to 2.2 across the eight human subjects, with a variation of 9.4%. The average technical variability of the same ratio from five replicate injections of a single sample was 17%. Again, using an authentic standard, we confirmed that tryptophan produced 4D features with a similar ratio. Next, we wanted to confirm that structural isomers had ratios across human subjects that were greater than those we observed for charge isomers. Given that we were unable to identify two co-eluting structural isomers in our dataset that were uniquely separated by IMS, we extended our search space to potential structural isomers that were separated by LC (i.e., 3D features that had the same *m/z* value but different retention times). One such example was *m/z* 274.0896 in positive mode, which appeared as two LC peaks at 4.0 and 6.8 min. Both 3D features were injective, meaning that each LC signal produced one IMS signal. The two LC peaks clearly had different ratios across the biological replicates, ranging from 6 to 62. The variation in the ratio of the LC peaks from the biological replicates was 66%, which was larger than the 14% variation that was observed between five technical replicates of the same sample.

Having established that our ratiometric approach can differentiate between types of non-injective 4D features, we focused on estimating the number of structural isomers in DTIMS data. Among the co-eluting non-injective 4D features that occurred in all eight subjects, most had relatively similar ratios that varied by less than 30%. In contrast, only a handful had clearly different ratios that varied by more than 40% (**Figure 3e**). Our conclusion from the analysis is that only a handful of non-injective 4D features represent structural isomers.

To further corroborate the claim that most non-injective 4D features are not structural isomers, even in our LC-TIMS-MS analysis, we moved to analyzing standards. Specifically, we evaluated our PolarMix comprised of 65 authentic chemical standards. From three independent runs, the average number of 3D features returned by XCiMS was 75, which included common degeneracies such as adducts. As with human plasma, the number of 4D features increased significantly to almost 150 (**Figure 3f**). Given that our PolarMix analysis did not contain structural isomers or isobars, these data demonstrate that nearly half of 3D features produce multiple 4D features due to factors such as charge isomers and ion clusters. We conclude that, while the IMS methods we applied do have the potential to resolve a small number of co-eluting chemicals, they increase the complexity of the dataset considerably. Most of the 4D features are a form of chemical degeneracy, which may provide limited biological insight.

### XCiMS shows that ion mobility reduces chimeric spectra by nearly half

We and others have shown that chimeric spectra are prevalent in metabolomics, even when standard chromatographic gradients are used with narrow MS/MS isolation windows.^39–41^ Chimeric spectra are problematic because they hinder metabolite identification. In contrast to MS/MS data in reference libraries, which are derived from the analysis of pure chemical standards, chimeric spectra contain fragments from multiple metabolite precursors. Consequently, chimeric spectra cannot be searched against reference libraries directly. Although several strategies have been established to deconvolve chimeric data^39, 40, 42^, they are not always effective and the metabolites contributing to the spectra can go unidentified.

A promising strategy to improve metabolite identification is to reduce the incidence of chimeric spectra in profiling experiments. While the benefits of IMS have most often been considered in the context of differentiating structural isomers, we sought to explore whether IMS separation of isobars might reduce the frequency of chimeric data (**Figure 4a**). To test this idea, we analyzed human plasma by LC-DTIMS-MS and processed the data with XCiMS. Next, using collapsed 3D data, we applied the same computational approach that we have described before. In brief, we counted features that would be co-isolated for fragmentation when using a 1.0 Da isolation window. In line with prior results^40^, we determined that 38% of 3D signals isolated for MS/MS would be chimeric. We then repeated the analysis, but this time considered the IMS dimension of the data. Strikingly, we found that IMS reduced the number of chimeric spectra by nearly half. When evaluating the 4D data, only 19% of features were chimeric (**Figure 4b**). The percentage of chimeric spectra was slightly higher in negative mode compared to positive mode. For those MS/MS spectra that did contain fragments from more than one precursor, we also estimated the extent to which they were chimeric. Beyond increasing the number of pure MS/MS spectra, we observed that IMS improved the overall purity of chimeric data (**Figure 4c-d**). The results demonstrate that IMS can substantially reduce chimeric spectra in standard metabolomics experiments and thus facilitate the process of metabolite identification.

**Figure 4.**
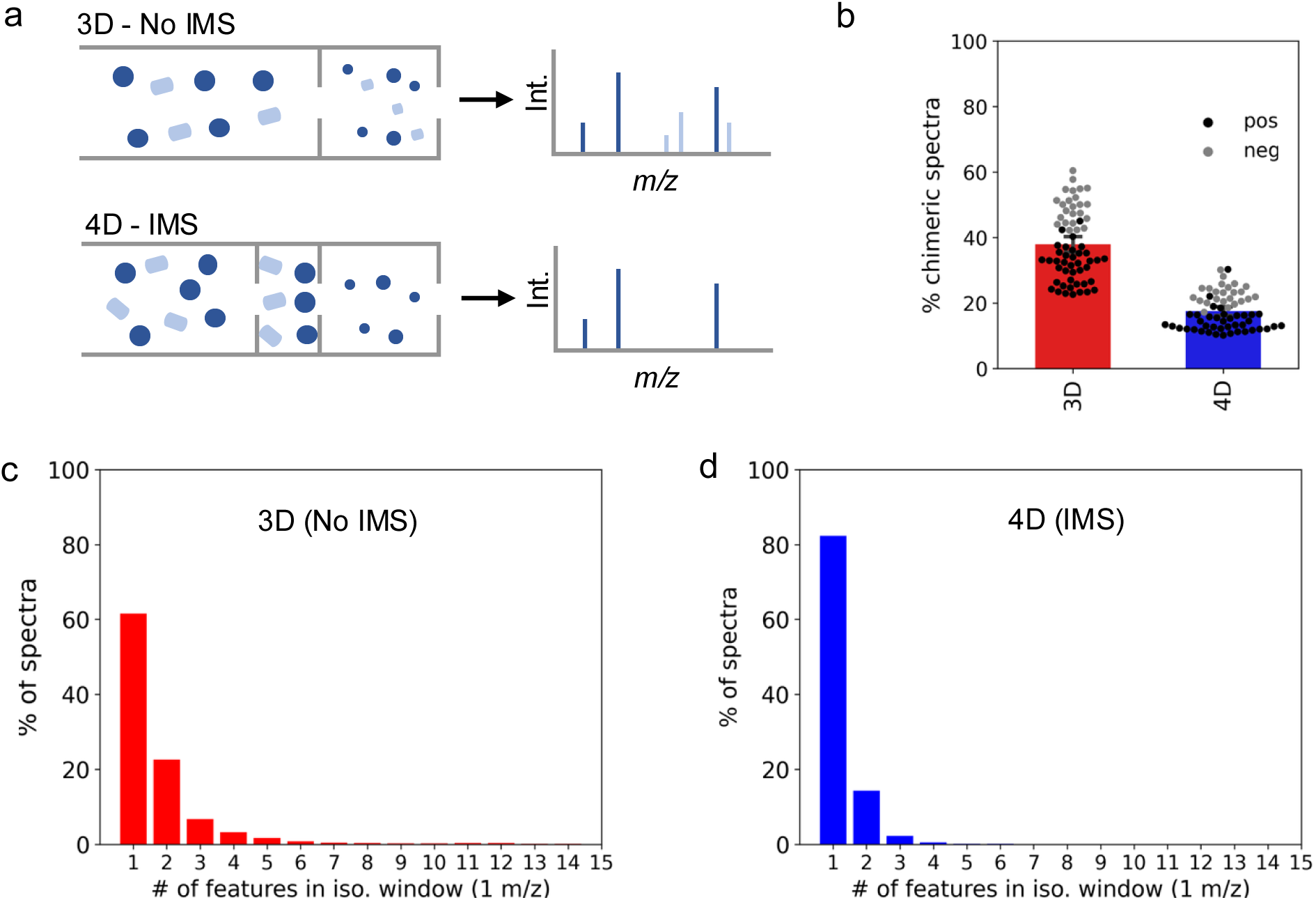
IMS reduces the frequency of chimeric spectra. a) Schematic to illustrate the origin of chimeric MS/MS spectra. Two co-isolated precursors, colored light and dark blue, are simultaneously fragmented and produce chimeric MS/MS spectra. The introduction of IMS separates the two precursors prior to fragmentation, preventing the generation of chimeric MS/MS spectra. b) Percentage of chimeric spectra obtained when performing untargeted metabolomics with and without IMS. The analysis is based on a 1 Da MS/MS isolation window. Data were acquired from human plasma by using a 16-minute HILIC method and DTIMS-MS. Individual dots represent biological (n=8) and technical (n=5) replicates from positive (black) and negative (grey) mode. c-d) IMS reduces the degree of contamination in chimeric spectra. When considering chimeric spectra (38% for 3D profiling and 19% for 4D profiling), IMS reduced the average number of contaminating precursors contributing to an MS/MS experiment. In other words, even though there were still chimeric spectra with the addition of IMS, the remaining chimeric spectra were less contaminated on average compared to chimeric spectra from 3D analyses.

## Discussion

XCMS is a widely used software package for processing untargeted metabolomics data from LC-MS experiments.^19, 20^ Here, we extend the platform to enable processing of 4D metabolomics data by establishing XCiMS. The first step in the XCiMS processing pipeline is the collapse of 4D data into pseudo-3D data. Peak detection is then performed in the pseudo-3D space to define regions of interest, from which targeted extraction of 4D signals occurs. We demonstrate that XCiMS is compatible with three prevalent ion mobility technologies: TWIMS, TIMS, and DTIMS from Waters, Bruker, and Agilent, respectively.

Similar to other data-processing software, including XCMS, the output of XCiMS is a table of features. Although structural identification requires matching experimental data to reference data from chemical standards, feature tables can provide important insight as a global readout of an experiment. Historically, for example, feature counts were used to estimate metabolite coverage across different analytical protocols and instruments.^25^ While this approach is not without limitations, it played a key role in guiding the development of untargeted metabolomics workflows.^26–30^ With the increasing availability of commercial IMS platforms, similar methodological challenges must now be addressed to establish effective 4D approaches for global profiling. As demonstrated by our analysis of TWIMS data, counts of credentialed features provide a practical metric for evaluating analytical parameters. In this context, just as XCMS helped catalyze the development of conventional metabolomics workflows, XCiMS is poised to accelerate the evolution of 4D metabolomics.

The processing architecture of XCiMS enables users to move back and forth between the 3D and 4D space of untargeted metabolomics data. This capability provides a unique opportunity to assess a longstanding question in the field: what is the benefit of incorporating ion mobility into untargeted metabolomics workflows? To facilitate the analysis, we found it helpful to classify 4D features as being injective or non-injective. When a 3D feature expands into a single 4D feature, we call it injective. Conversely, when a 3D feature expands into multiple 4D features, we refer to them as being non-injective. IMS holds promise for differentiating structural isomers that cannot be resolved by LC-MS. Such cases would be expected to produce non-injective 4D features. Indeed, XCiMS revealed that IMS can lead to thousands of non-injective 4D features, depending on the technology being used. Notably, however, a deeper investigation into the origin of the non-injective 4D features demonstrated that most are not a result of structural isomers. Instead, the majority of non-injective 4D features were caused by a combination of charge isomers and ion clusters.

We find it useful to draw an analogy between charge isomers and adducts. Both represent a form of degeneracy, causing one chemical to appear in the data as multiple signals. Like adducts, charge isomers increase the complexity of untargeted metabolomics data without adding biological information. For adducts, numerous software solutions are available to cluster features representing the same metabolite.^35, 43–45^ In positive mode, for instance, it is relatively routine to group protonated and sodiated signals derived from the same parent compound. At present, there is not a comparable computational tool for annotating charge isomers from untargeted metabolomics data. Here, we estimated the number of charge isomers by identifying pairs of non-injective 4D features that exhibited consistent intensity ratios across all human subjects analyzed by the same experimental methods. Moving forward, it will be helpful to establish additional annotation approaches for charge isomers that can be applied at the level of individual samples. Without such a development, our ability to interpret metabolomics data from 4D experiments may be limited.

Given the considerable increase in data complexity introduced by IMS, largely through signal degeneracy, its incorporation into untargeted metabolomics workflows requires a compelling analytical justification. Although the ability to differentiate structural isomers is often assumed to be a key rationale for IMS, our results suggest otherwise. We show that the addition of IMS does not markedly improve resolution of structural isomers, at least when using standard chromatographic gradients applied to human plasma. Hence, this capability alone does not provide sufficient rationale for performing 4D profiling. Rather, our findings indicate that the primary value of IMS in untargeted metabolomics lies in improving the quality of MS/MS data through reduction of chimeric spectra.

Although chimeric spectra are commonly associated with the wide MS/MS isolation windows used in data-independent acquisition, they are also prevalent in metabolomics workflows employing narrow precursor selection.^40^ Chimeric spectra complicate compound identification because they generally do not match reference spectra acquired from pure chemical standards, which form the basis of most spectral libraries. Over the past decade or so, several computational strategies have been established to deconvolve chimeric spectra, but they are not successful in every case. ^39, 40, 42^ To improve compound identification rates, we therefore need to minimize the simultaneous fragmentation of co-eluting precursors. Our results demonstrate that IMS is highly effective in this regard, reducing the frequency of chimeric spectra by approximately half when human plasma is analyzed with a standard chromatographic gradient and a 1 *m/z* MS/MS isolation window. Collectively, these findings suggest that the greatest impact of IMS in untargeted metabolomics may come not from increasing the number of detected features, but from increasing the reliability with which features can be translated into chemical structures.

## Methods

### Chemicals and reference materials

LC-MS grade acetonitrile (ACN), water, isopropanol (IPA), methanol (MeOH), ammonium bicarbonate, ammonium formate, ammonium hydroxide, and methylenediphosphonic acid (medronic acid) were purchased from MilliporeSigma. Amino acid standards were purchased from MilliporeSigma. The Mass Spectrometry Metabolite Library of Standards (MSMLS) was purchased from IROA Technologies. Compounds for the polar metabolite standard mixture (PolarMix) were purchased from MilliporeSigma. Standard mixtures were prepared in ACN/water (1:1, v/v). Amino acid standards were prepared at a final concentration of 10 µM. The MSMLS standard mixture was prepared according to instructions on the product sheet followed by pooling of standards from plate 1-6. PolarMix was made by preparing stock solutions for individual standards at concentrations of 10 or 100 mM and mixing to a diluted concentration of 10 µM. The plasma reference material SRM1950 was purchased from the National Institute of Standards and Technology (NIST). The credentialed E. *coli* sample was prepared by using the Credentialed E. *coli* Cell Extract Kit purchased from Cambridge Isotope Laboratories (MSK-CRED-DD-KIT). Briefly, ^12^C-unlabeled and ^13^C-labeled E. *coli* extracts in the kit were reconstituted in 100 µL of ACN/water (1:1, v/v), vortexed for 30 seconds, sonicated for 5 minutes, and centrifuged at 14,000 rpm and 4 °C for 10 minutes. The supernatants were transferred to LC-MS vials separately. ^12^C and ^13^C samples were mixed at 1:1 and 1:2 ratios (v/v) in two LC-MS vials as the credentialed samples (“1T1” vs. “1T2”).

### Human plasma samples

Plasma from eight individuals was obtained from the Nutrition Obesity Research Center at Washington University in St. Louis. The use of plasma from the participants was approved by the Institutional Review Board within the Human Research Protection Office at Washington University School of Medicine in St. Louis. All participants provided written, informed consent before plasma samples were collected.

#### Metabolite extraction of human plasma

Polar metabolites from NIST SRM1950 plasma and plasma from human subjects at Washington University in St. Louis were extracted by adding 450 µL MeOH to 50 µL plasma. The samples were vortexed two times for 30 seconds. The extracts were then placed at -20 °C overnight for protein precipitation before being centrifuged at 14,000 rpm and 4 °C for 15 minutes. The supernatants were transferred to LC-MS vials.

#### LC-TWIMS-MS analysis

The analysis was performed by using a Waters Acquity I-Class UPLC system interfaced to a Waters SYNAPT XS mass spectrometer equipped with traveling wave ion mobility spectrometry (Waters Corporation). An aliquot of 2 µL of sample was injected per analysis. A 100 mm iHILIC-(P) Classic column coupled to a 20 mm guard column (2.1 mm, 5 µm, 200 Å, HILICON) was used. Mobile-phase A consisted of 5:95 ACN/water (v/v) with 20 mM ammonium bicarbonate, 0.1% ammonium hydroxide (v/v), and 2.5 µM medronic acid. Mobile phase B consisted of 95:5 ACN/water (v/v) and 2.5 µM medronic acid. The column was operated at 40 °C with the following gradient and flow rate: 0 min, 90% B at 0.25 mL/min; 1 min, 90% B at 0.25 mL/min; 12 min, 35% B at 0.25 mL/min; 12.5 min, 25% B at 0.25 mL/min; 14.5 min, 25% B at 0.25 mL/min; 15 min, 90% B at 0.25 mL/min; 15.5 min, 90% B at 0.25 mL/min; 16.5 min, 90% B at 0.4 mL/min; 20 min, 90% B at 0.4 mL/min; 20.5 min, 90% B at 0.25 mL/min; 22 min, 90% B at 0.25 mL/min. Unless otherwise specified, the TWIMS-MS instrument was operated with the following parameters: Capillary Voltage, 2.2 kV for negative mode; Source Temperature, 140 °C; Desolvation Temperature, 300 °C; Desolvation Gas Flow, 1000 L/hour; Nebulizer, 6 bar; IMS Wave Velocity, 1000-250 m/s with 100% Ramp Time; IMS Wave Height, 31 V; IMS Nitrogen Gas Flow, 90 mL/min; TOF Mode, Resolution Mode with 66 µs ADC Pusher Frequency; Mass Range, 50-1200 m/z; Scan Duration, 1 sec; EDC Delay Coefficient, 1.41 for negative mode. CCS calibration was performed by establishing the calibration equation using the Waters Major Mix Calibration Kit with calibrant ions of known CCS values.

#### LC-DTIMS-MS analysis

Human plasma extracts were separated by HILIC as described above, using a 100 mm iHILIC-(P) Classic column coupled to a 20 mm guard column (2.1 mm, 5 µm, 200 Å, HILICON). Mobile-phase A consisted of 5:95 ACN/water (v/v) with 20 mM ammonium bicarbonate, 0.1% ammonium hydroxide (v/v), and 2.5 µM medronic acid. Mobile phase B consisted of 95:5 ACN/water (v/v). The column was operated at 40 °C with the same gradient and flow rate as described for LC-TWIMS-MS. The separation was carried out by using an Agilent 1290 Infinity II UHPLC system. The injection volume was 1 μL. The drift tube ion mobility spectrometer utilized for IMS-MS was an Agilent 6560 system. The instrument consists of a 78 cm uniform field drift tube pressurized with high purity nitrogen drift gas. Analytes were ionized by using the Agilent Jet Stream electrospray source in positive or negative mode (3500 V and 3000 V, respectively). The following parameters were applied: gas temperature 300°C, gas flow 10 L/min, nebulizer 35 psi, sheath gas temperature 350°C, and sheath gas flow 11 L/min. In accordance with the standardized protocol described previously^46^, the pressure in the drift tube was maintained at 3.95 Torr and the applied separation voltage was 1574 V.

#### LC-TIMS-MS analysis

The analysis was performed with an Agilent 1260 HPLC system interfaced with a Bruker timsTOF fleX MALDI-2 mass spectrometer. An aliquot of 2 µL of sample was injected for each analysis. The LC method used in this analysis was the same as for the LC-TWIMS-MS analysis. The mass spectrometer was operated under following settings: dry gas, 10 L/min; dry gas temperature, 220°C; nebulizer, 2.5 bar; capillary voltage, 3.6 kV; end plate offset, 500 V; TIMS 1/K_0_ scan range, 0.1 - 1.4 Vs/cm^2^; TIMS accumulation time: 250 ms; TIMS ramp time: 250 ms; TIMS ion charge control: 7.50 M; and mass range, 50-1050 *m/z*. TIMS and MS calibration was performed by using a mixture of Agilent ESI-L Tuning Mix and 10 mM sodium formate calibrant solution (9:1, v/v) at the beginning of each experiment. The calibrant was also infused in between 0.1 and 0.3 min of each analysis for internal calibration.

#### XCiMS workflow

An overview of the entire workflow is shown in **Supplementary Figure 8**. Each section of the workflow is described separately below.

##### Data Input

The XCiMS workflow starts by parsing MS raw data files into an *LCIMMSpectraL* object, which includes vendor-agnostic LC-IMS-MS spectra. To reduce memory cost, only spectra ID, *m/z*, RT frame, IM bin, and intensities are extracted. XCiMS can parse multiple .mzML files by using the *constructLCIMMSpectraL* function. This function utilizes the *MSnbase* package to access .mzML data. By using software such as ProteoWizard’s MSConvert^47^, most vendor-proprietary files can be converted to .mzML files including Agilent .D for TIMS and Waters .raw for TWIMS. At the time of doing the analysis, however, it was not practical to convert Bruker .d files with this approach because the size of the .mzML files commonly exceeds 30 GB per sample, which is about 100 times bigger than the original file. To address this issue, we developed an alternative function, *constructLCIMMSpectra2L*, to parse Bruker .d files directly. This is achieved by leveraging a recently developed R package called *opentimsr*. We note that *opentimsr* only works in Windows and Linux systems and requires the user to accept Bruker’s license agreement. Therefore, processing of TIMS data is limited to Windows or Linux operating systems.

##### Generate pseudo LC-MS data by spectral binning

The second step of the workflow is to generate a *pLCMSpectraL* object from the *LCIMMSpectraL* object by using the *constructpLCMSpectraL* function. This step creates pseudo LC-MS spectra of which the ion mobility (IM) dimension is collapsed. To achieve this, an efficient binning algorithm was developed that can combine any number of MS spectra into one spectrum (**Supplementary Figure 1b**). Here, we detailed the procedure to generate a pseudo LC-MS spectrum using the binning algorithm. To start, we denote a four-dimensional (4D) MS spectrum with *n* peaks at RT frame

*x* and IM bin *y* as in **Equation 1**:

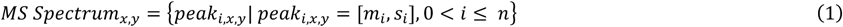

where [*m_i_*, *s_i_*] represents *m/z* and intensity pair for peak *i*.

Suppose the LC-IMS-MS data has *a* RT frames and *b* IM bins, the corresponding pseudo LC-MS spectra are generated in 6 steps:

1. A peak set with *n m/z* (*M_k_*) and intensity (*S_k_*) pairs, *P_k_*, is extracted from all MS spectra at RT frame *k* as in **Equation 2**:

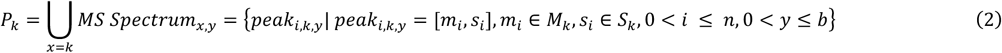
2. An ordered peak set, *O_k_*, is initiated by sorting all *m/z* values in ascending order from *P_k_* as in Equation 3:

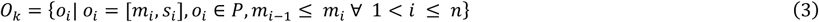
3. A numeric array, *A_k_*, is generated by calculating relative differences of adjacent *m/z* centroids in *O_k_* as in **Equation 4**:

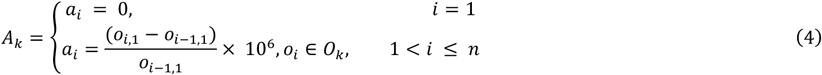

A binary array, *B_k_*, is generated to indicate whether the relative differences denoted in *A_k_* is larger than a threshold value, *t*
(*t* = 0.1 by default) as in **Equation 5**:

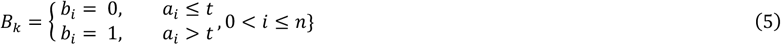
4. A splitting array, *C_k_*, is generated by calculating the cumulative sum of *B_k_* as in **Equation 6**:

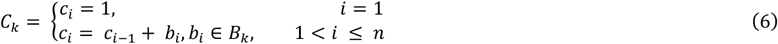

Let *U_k_* be a set of all unique values in *C_k_*. A collection, *D_k_*, composed of |*U_k_*| subsets of *O_k_* is constructed as in **Equation 7**:

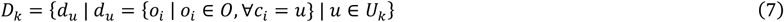
5. A new peak set, *Q_k_*, is generated by calculating the intensity weighted mean of each subset in *D*, as in **Equation 8**, where *Q_k_* is assigned as the pseudo LC-MS spectrum at RT frame *k*:

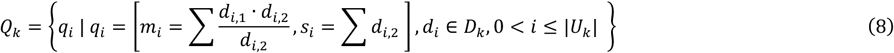
6. Step 1-5 is repeated for each RT frame *x* to generate pseudo LC-MS spectra of one data file as in Equation 9:

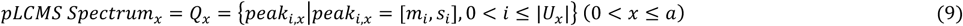

##### Multidimensional feature detection

XCiMS performs multidimensional feature detection by exploiting the *centWave* algorithm in the XCMS ecosystem in three steps:

1) Three-dimensional (3D) precursors, conventionally known as LC-MS features, are detected from the pseudo LC-MS data by using the *centWave* algorithm in the XCMS R package as previously described.^21^ Briefly, in pseudo LC-MS data, continuous ion traces along the RT dimension within a small *m/z* window are detected as regions-of-interest (ROIs). Chromatographic peak detection on each ion trace (i.e., an extracted ion chromatogram or EIC), is performed by using the continuous wavelet transform (CWT) algorithm. Collectively, 3D precursors (pseudo LC-MS features) are exported with an *m/z* window (*mz*, *mzmin*, *mzmax*), chromatographic peak profile (*rt*, *rtmin*, *rtmax*), peak area and height (*into, maxo*), signal-to-noise ratio (*sn*), and baseline corrected-peak area (*intb*).^19^ A unique *index* number is assigned to each 3D precursor as the ID. In XCiMS, 3D precursors are detected from the *pLCMSpectraL* object and exported in a *pLCMSpeaks* object by using the *findPeaks_centWave_pLCMSL function*.
2) Using the *m/z* and chromatographic peak profiles of each 3D precursor, a region-of-interest in the RT-IM dimension (RT-IM ROI) is extracted, where the intensities of all *m/z* peaks are extracted and summed from LC-IMS-MS spectra. The same *index* is assigned to each RT-IM ROIs as an ID corresponding to the 3D precursor. The RT-IM-ROIs are extracted from the *LCIMMSpectraL* object in step 1 and exported in a *RtImMapL_pLCMSpeaks* object by using the *extratRtImMapL_pLCMSpeaks* function with the *pLCMSpeaks* object in step 1.
3) An ion trace along the IM dimension (i.e., the extracted ion mobilogram or EIM) is extracted for each RT-IM ROI by summing the intensities of all data points at each IM bin. Then, IMS peak detection on each EIM is performed by using the CWT algorithm built-in a tidy variant of the *centWave* algorithm in the XCMS package.^47^ IMS peak profiles are reported with *index* (ID of the RT-IM-ROI), *im, immin*, *immax*, and *intn* (the sum of all intensities in between *immin* and *immax*). A unique 4D feature is generated by joining each IMS peak to its corresponding 3D precursor through the *index* value, along with a unique *index4d* number assigned to it as a new ID. 4D features are detected from the *RtImMapL_pLCMSpeaks* object and exported in an *LCIMMSpeaks* object by using the *findPeaks_centWave_MobilL* function with the *pLCMSpeaks* object in step 2.

##### Multidimensional feature correspondence

XCiMS is developed with the capability to process multiple data files. Using LC and IMS peak profiles, XCiMS can perform 4D feature correspondence by using the *density* algorithm within the XCMS ecosystem to organize 4D features into feature groups.^19^ In LC-MS data, feature groups are determined by splitting features to small *m/z* slices, estimating the density distribution of all LC peaks at each *m/z* slice, and grouping features at dense regions with similar retention times.^19^ Similarly, in LC-IMS-MS data, feature correspondence starts with grouping 4D features based on LC peak density and further organizes them using IMS peak information and the *density* algorithm with a smaller standard deviation setting.

In XCiMS, 4D features in the *LCIMMSpeaks* object are grouped by using the *groupPeaks_density_LCIMMS* function to generate an *LCIMMSpeakgroups* object. We note that XCiMS also supports feature correspondence for pseudo LC-MS data (*groupPeaks_density_pLCMS*) and pseudo IM-MS data (*groupPeaks_density_pIMMS*).

##### Fold-change analysis

When two classes of samples are analyzed, XCiMS performs fold-change analysis by performing a Welch t-test among the intensities (peak areas) of 4D features in each multidimensional “meta-peak” and calculating the ratio of mean intensities between two classes. Fold-change analysis is performed on the *LCIMMSpeakgroup* object by using the *FCAnalysis_LCIMMSpeaks* function, with the result being exported in a *statsummary* object. We note that fold-change analysis can also be performed on pseudo LC-MS feature groups and pseudo IMS-MS feature groups by using the *FCAnalysis_pLCMSpeaks and FCAnalysis_pIMMSpeaks* functions.

##### CCS calculation

For each type of IMS data supported by XCiMS, an individual function is built into the XCiMS R package for CCS calculation:

*CalCCSTWIMS* function: The unit of the IMS dimension in TWIMS data is drift time in milliseconds. CCS calibration is required to accurately calculate CCS values. Each TWIMS CCS calibration establishes a fitted equation with three parameters *t*_#_, exponent *m*, and coefficient *k*. These parameters are used to calculate TWIMS CCS values as in **Equation 11**:

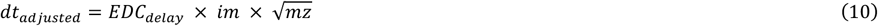

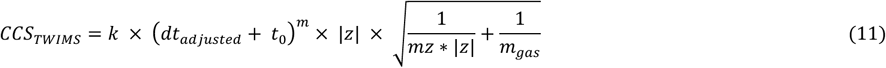

where *EDC_delay_*, *m*_)*as*_, *im*, *mz*, *z* are the EDC delay coefficient of the TWIMS-MS instrument, mass of the IMS buffer gas (generally N2, 28.0062), IM peak apex, mass-to-charge value, and charge state (generally 1 or -1) of 4D features or pseudo IMS-MS features, respectively.

*CalCCSTIMS* function: The unit of the IMS dimension in TIMS data is inverse reduced mobility 1/*K*_#_ in Vs/cm^2^. CCS calibration is not required for calculation, but it is required to measure 1/*K*_#_ values accurately. TIMS CCS values are calculated by using an empirical variant of the Mason-Schamp equation as in **Equation 12**:

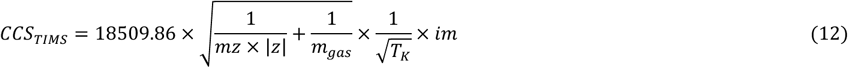

where *T_K_*, *m*_)*as*_, *im*, *mz*, *z* are room temperature in Kelvin, mass of IMS buffer gas (generally N2, 28.0062), IM peak apex, mass-to-charge value, and charge state (generally 1 or -1) of 4D features or pseudo IMS-MS features, respectively.

*CalCCSDTIMS* function: The unit of the IMS dimension in DTIMS data is drift time in milliseconds. CCS calibration is required to calculate CCS values accurately from a DTIMS system operated with a single-field method. DTIMS CCS calibration will provide two parameters *t_fix_*, and *β*, which are then used to calculate DTIMS CCS values as in **Equation 13**:

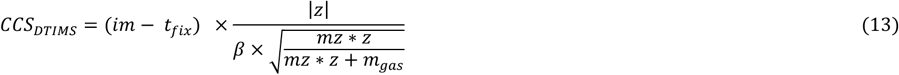

where *m*_)*as*_, *im*, *mz*, *z* are the mass of IMS buffer gas (generally N_2_, 28.0062), IM peak apex, mass-to-charge value, and charge state (generally 1 or -1) of 4D features or pseudo IMS-MS features, respectively.
**Supplementary Figure 9** shows the correlation between XCiMS-calculated CCS values and CCS values from the Unified CCS Compendium of polar metabolite standards measured via LC-TWIMS-MS and LC-DTIMS-MS with a mean error of 1.4%.

We note that the functions and objects in XCiMS suffixed with “*X*” are for single file processing, XCiMS objects and functions suffixed with “*L*” are for multi-file processing. For simplicity, we use objects and functions suffixed with “*L*” to explain the workflow. We note that a similar function *constructpIMMSpectraL* is also developed to construct pseudo IM-MS spectra for processing flow injection-IMS-MS (FI-IMS-MS) or direct infusion-IMS-MS (DI-IMS-MS) data. A flowchart on how to process LC-IMS-MS data and FI-IMS-MS data step-by-step by using functions in XCiMS is provided in **Supplementary Figure 10**. We recommend running XCiMS on a computer with at least 32 GB RAM, especially when multi-file processing is needed.

#### Optimization of *centWave* parameters for IMS peak detection

The separation mechanisms and units of the IMS dimension are different among TWIMS, DTIMS, and TIMS instruments, resulting in distinct IMS peak profiles. To optimize *centWave* parameters, the percentage of 3D precursors with detected IMS peaks is used as a metric to evaluate the performance of IMS peak detection. Since total 3D precursors could include informatic artifacts, we refer to 3D precursors of authentic standards, either from the amino acid standard mixture (AAMix) or the polar metabolite standard mixture (PolarMix) as the reference signals. *centWave* parameters including *peakwidth*, *prefilter*, *snthresh*, and *noise* were tuned to maximize IMS peak detection of reference signals.

#### XCiMS parameters

Unless otherwise specified, the XCiMS parameters described here were used to process all datasets:

- The following parameters were used in the *findPeaks_centWave_pLCMSL* function for chromatographic peak detection on pseudo LC-MS spectra: *ppm=20*, *peakwidth = c(10, 60), prefilter = c(3, 500), snthresh=5, noise=100*
- The following parameters were used in the *findPeaks_centWave_MobilL* function for IMS peak detection by IMS type:

- TWIMS datasets: *peakwidth = c(0.5, 2), prefilter = c(3, 500), snthresh=5, noise=0*
- DTIMS datasets: *peakwidth = c(0.4, 4), prefilter = c(3, 500), snthresh=5, noise=0*
- TIMS datasets: *peakwidth = c(0.02, 0.2), prefilter = c(3, 500), snthresh=5, noise=100*
- The following parameters were used in the *extractRtImMapL_pLCMSpeaks* function to extract RT-IM ROIs for all datasets: *ppm=25, RTscanExtend=0.1, mzmode=”centroid”*
- The following parameters were used in the *groupPeaks_centWave_MobilL* function for multidimensional feature correspondence:

- Chromatographic grouping: *bw=3, minSamples=3, binSize=0.025*
- IMS grouping: *bw=0.1, minSamples=3, binSize=0.025*
- The following parameters were used for CCS calculations by IMS type:

- *CalCCSTWIMS: EDCdelay=1.41, GasMass=28.0062, CCScoeff=140.68, CCSexp=0.979463, CCSt0=1.56822*
- *CalCCSDTIMS: Beta=0.1506, DTfix=0.4033, GasMass=28.0062*
- *CalCCSTIMS: gasMW=28.0062, roomTemperature_c=25*

Our HILIC gradient consists of an analytical segment (0-16 minutes) and an equilibration segment (16-22 minutes). Therefore, we only extracted and processed 4D MS spectra from the first 16 minutes. This is achieved by setting parameter *rtrange=c(0,960)*in the *constructLCIMMSpectraL*and *constructLCIMMSpectra2L* functions.

### Using XCiMS to optimize LC-IMS-MS platforms for untargeted metabolomics

XCiMS can unbiasedly detect multidimensional features from the raw data and can thus be used to optimize LC-IMS-MS platforms for untargeted metabolomics systematically. Using the TWIMS-MS system as an example, we evaluated the performance of the instrument for untargeted profiling under three parameter sets (**Supplementary Figure 4a**). We first evaluated two sets of wave heights and wave velocities under TOF resolution mode (parameter set 1 and parameter set 2) using XCiMS and the *credentialing* workflow we previously developed (**Supplementary Figure 4b)**.^25^ Metabolite extracts of unlabeled and fully ^13^C-labeled were mixed at two specific ratios (1:1 and 1:2) to create a pair of credentialed reference samples. The samples were analyzed by LC-TWIMS-MS under two IMS parameters settings. The data were processed by XCiMS to detect 3D precursors and 4D features followed by applying the *credential* R package to search for ^12^C and ^13^C counterparts that pass the intensity ratios in the feature table. These biologically relevant features represent the unlabeled and labeled versions of metabolites synthesized de novo by E. *coli*.

We also tested two TOF acquisition modes: high-resolution mode (parameter set 3) and resolution mode (parameter set 2). In principle, ions will travel faster with larger wave height or smaller wave velocity, both of which induce higher electric forces applied to the ion. As a result, the peaks would be narrower, requiring a faster scan rate by the TOF to ensure data sampling. Under high-resolution mode, XCiMS detected 3,642 3D precursors and 3,094 4D features from data acquired using parameter 3. In contrast, 17,144 3D precursors and 20,203 4D features were detected under resolution mode, increased by 470% and 660% (**Supplementary Figure 4c**). When TWIMS was operated at a ramped wave velocity from 1000 – 250 m/s and a wave height of 31V (parameter set 2), IMS peak widths of most 4D features fall in the range between 0.2 to 0.5 milliseconds. With high-resolution mode, <4 data points could be obtained to construct an ion trace along the IM dimension for those 4D features. In contrast, up to 8 data points were obtained under resolution mode with a TOF pusher frequency of 0.066 milliseconds per transition.

### Calculating the frequency of chimeric spectra

The frequency at which chimeric spectra occur in our data was calculated from our XCiMS feature table of the LC-DTIMS-MS data in positive and negative mode. A Top 3 DDA (1.0 m/z isolation window) LC/IMS/MS/MS experiment was simulated based on the XCiMS feature table. This simulation is an iterative process that traverses the RT and IM space and examines the number of co-isolated features. First, 1000 RT frames were created that were linearly spaced between the minimum and maximum RT values for the dataset (in this case 5.3 and 957.34 seconds). Next, in each RT frame, 100 IM frames were created that spanned the minimum and maximum IM values (in this case 0.08 ms and 59.03 ms). The process began at the first RT frame and iterated over the 100 IM frames. For each IM frame, we selected up to three targets for MS/MS fragmentation based on the three most intense features from the XCiMS peak table that have a RT range, [rtmin, rtmax], overlapping the RT frame and that have an IM range, [immin, immax], overlapping the IM frame. Then, for each target feature, we gathered all features from the XCiMS peak table that also have RT and IM ranges that overlap the RT and IM frame and have an *m/z* value within 0.5 *m/z* of the target *m/z* value. When the only feature that meets these criteria was the target feature, we denoted it as a non-chimeric spectrum. Otherwise, we recorded the number of co-isolated features and classified the MS/MS scan as chimeric. This process was repeated for all RT and IM frames.

#### Data analysis using Skyline software

Targeted analysis on LC-TWIMS-MS data of AAMix samples and the spike-in experiment was performed by using the Skyline software (v21.2.0) with the following Transition Settings: *Precursor mass = “monoisotopic”, Ion match tolerance = 0.01 m/z; Precursor mass analyzer = “TOF”; Resolving power = 30,000*. The transition lists of AAMix and PolarMix were created based on the in-house RT library. The [M-H]^-^ion was used for quantitative analysis. Chromatographic peak areas of all standards were carefully integrated by manual inspection of the EICs.

#### Data analysis using Compass DataAnalysis software

Targeted analysis on LC-TIMS-MS data of PolarMix samples was performed by using Bruker Compass DataAnalysis software (v6.0). The EICs of [M-H] ions for all PolarMix standards were extracted with an *m/z* width of 0.01 Da. The average MS spectrum and EIM of each standard were extracted at corresponding chromatographic peak regions.

#### Data analysis using Agilent IM-MS Browser

EIMs for phenylalanine and tryptophan were obtained from IM-MS Browser 10.0 (Agilent Technologies).

## Acknowledgement

This work was supported by R35ES028365 (G.J.P.) from the NIH. We thank Aiko Barsch, Matthew Lewis, and Heiko Neuweger (Bruker Corporation) for their insights on analyzing charge isomers with TIMS. We thank Sheher Mohsin and John Sausen (Agilent Technologies) for their assistance in acquiring LC-DTIMS-MS data, Dr. Gordon Smith for providing the human plasma samples, and Dr. Steve Doonan for discussions related to LC-TWIMS-MS and LC-TIMS-MS analysis.

## Author contributions

L.W. and G.J.P designed the research. L.W., J.Y., and K.C developed the computational program and scripts. L.W., J.Y., and D.S.A performed experiments using LC-TWIMS-MS and LC-TIMS-MS analysis. L.W., J.Y., E.S., K.C., M.S.H, C.S.H, and T.G. performed data processing. J.N.D. and E.S.B. contributed to conceptualization and data interpretation. M.S.H. and K.C. prepared standard mixtures, created in-house libraries of metabolite standards, and performed data analysis. L.W., E.S, M.S.H. and G.J.P. wrote the manuscript. All authors contributed to revisions and approved the final version of the manuscript.

