## Supplementary Figures for "Using XCiMS to Quantify the Benefits of Ion Mobility Spectrometry for Untargeted Metabolomics"

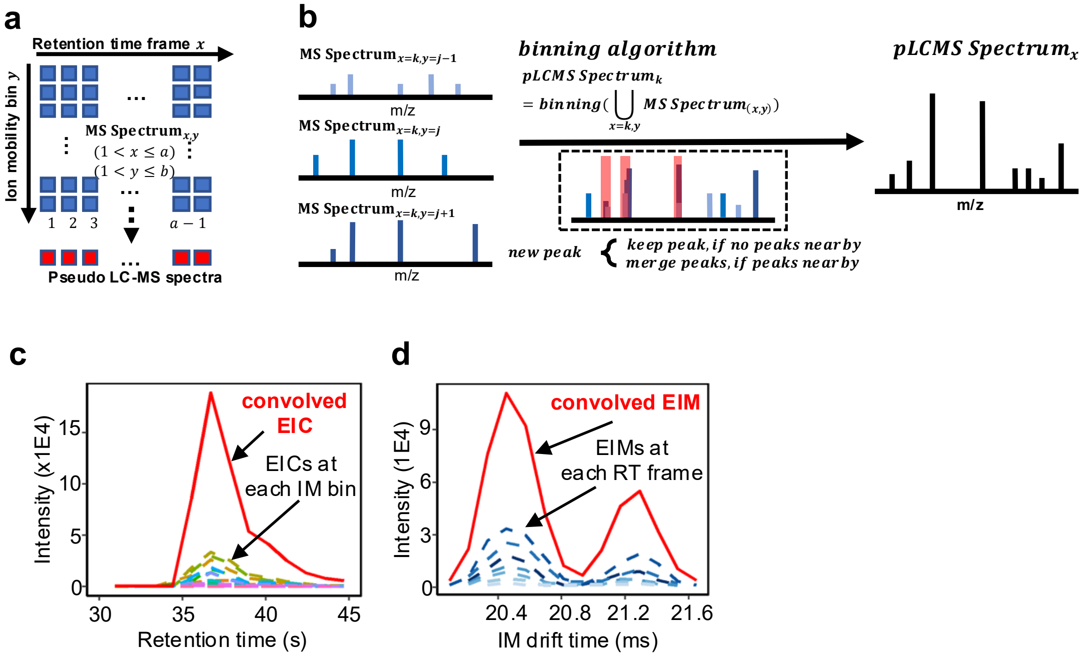


**Supplementary Figure 1. The XCiMS workflow.** a) Data from 4D MS spectra are partitioned into a matrix of RT frames and IM bins. b) A spectral-binning algorithm combines 4D MS spectra into a pseudo LC-MS spectrum. Three spectra are shown here for demonstrative purposes. c) Representative example of a convolved EIC (red solid line) versus discrete EICs extracted along each IM bin (dashed lines). d) Representative example of a convolved EIM (red solid line) versus discrete EIMs extracted along each RT frame (dashed lines).

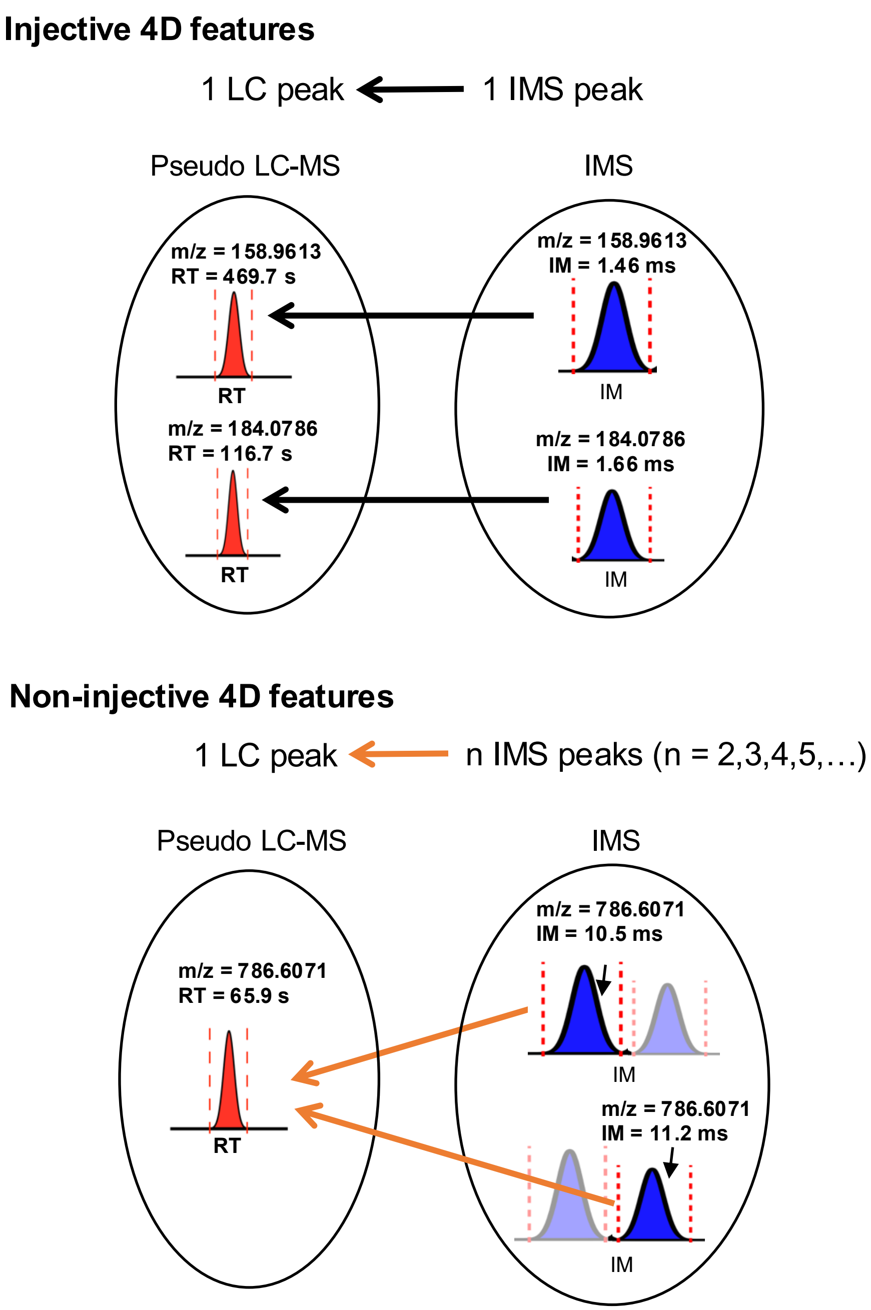


**Supplementary Figure 2. Injective and non-injective 4D features.** An injective 4D feature is a single 4D feature derived from a single 3D feature. Non-injective 4D features are multiple 4D features derived from the same 3D feature. A representative example of each is schematized here for demonstration.

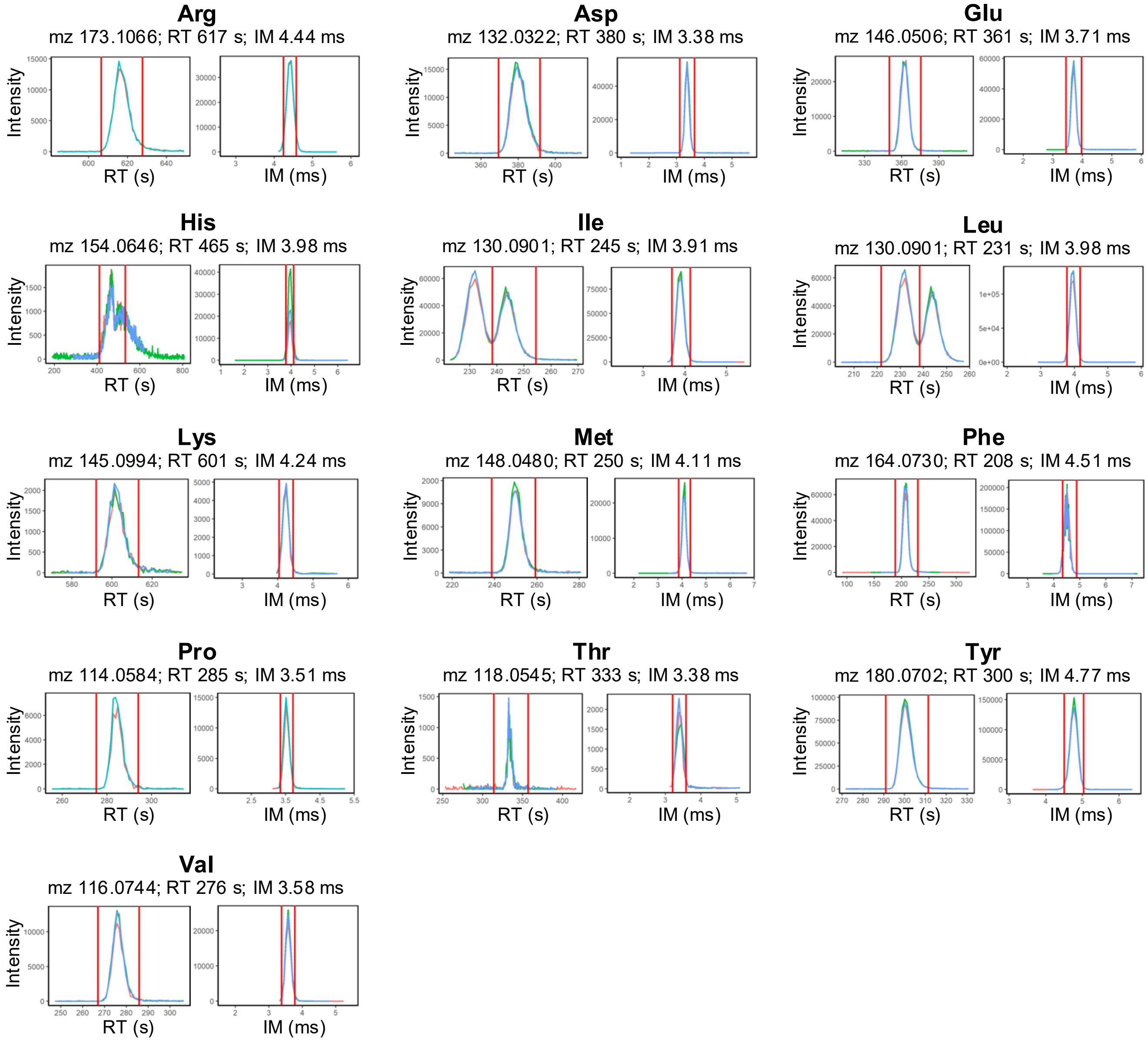


**Supplementary Figure 3. XCiMS performs multidimensional feature detection.** Extracted ion chromatograms (EIC) and extracted ion mobilograms (EIM) output from XCiMS for 13 amino acids analyzed in triplicates by using HILIC-TWIMS-MS analysis. Multidimensional feature detection and correspondence were performed by using the XCiMS software. LC and IMS peak profiles are shown with red lines indicating boundaries determined by XCiMS. The optimized parameters used for peak detection were *peakwidth = c(0.3, 0.9)*, *snthresh = 1*, *noise*, *prefilter = c(3, 500), noise = 100.*

**a**

**TWIMS-MS parameters for 4D untargeted metabolomics**

| | |
|---|---|
| TWIMS-MS parameter 1 | wave height: 25V;<br>wave velocity 1000-300 m/s;<br>Resolution mode (TOF pusher frequency: 66 us) |
| TWIMS-MS parameter 2 | wave height: 31V;<br>wave velocity 1000-250 m/s;<br>Resolution mode (TOF pusher frequency: 66 us) |
| TWIMS-MS parameter 3 | wave height: 25V;<br>wave velocity 1000-300 m/s;<br>High-resolution mode (TOF pusher frequency: 145 us) |

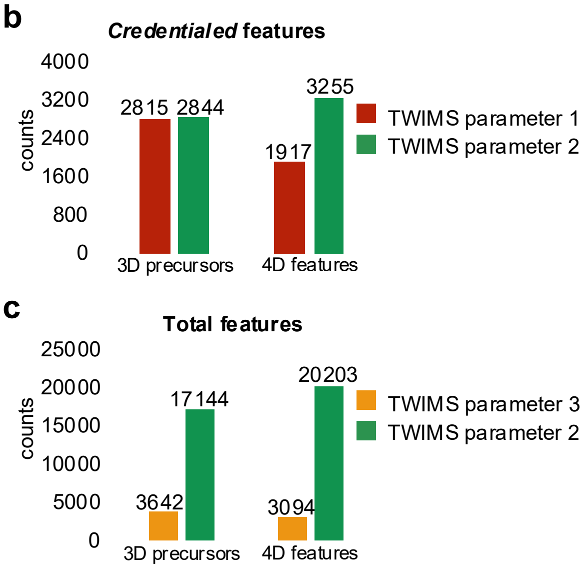


**Supplementary Figure 4. Optimizing traveling wave ion mobility spectrometry-mass spectrometry (TWIMS-MS) for 4D untargeted metabolomics.** a) A summary of the three sets of TWIMS-MS parameters evaluated. b) Number of credentialed 3D precursors and 4D features detected from *E. coli* samples by using the first and the second set of TWIMS-MS parameters. c) Number of total 3D precursors and 4D features detected from SRM1950 plasma extracts when using the second and third set of TWIMS-MS parameters. 4D features were detected by XCiMS and, for *E. coli*, filtered based on isotopic signatures with a standard credentialing workflow (see Methods).

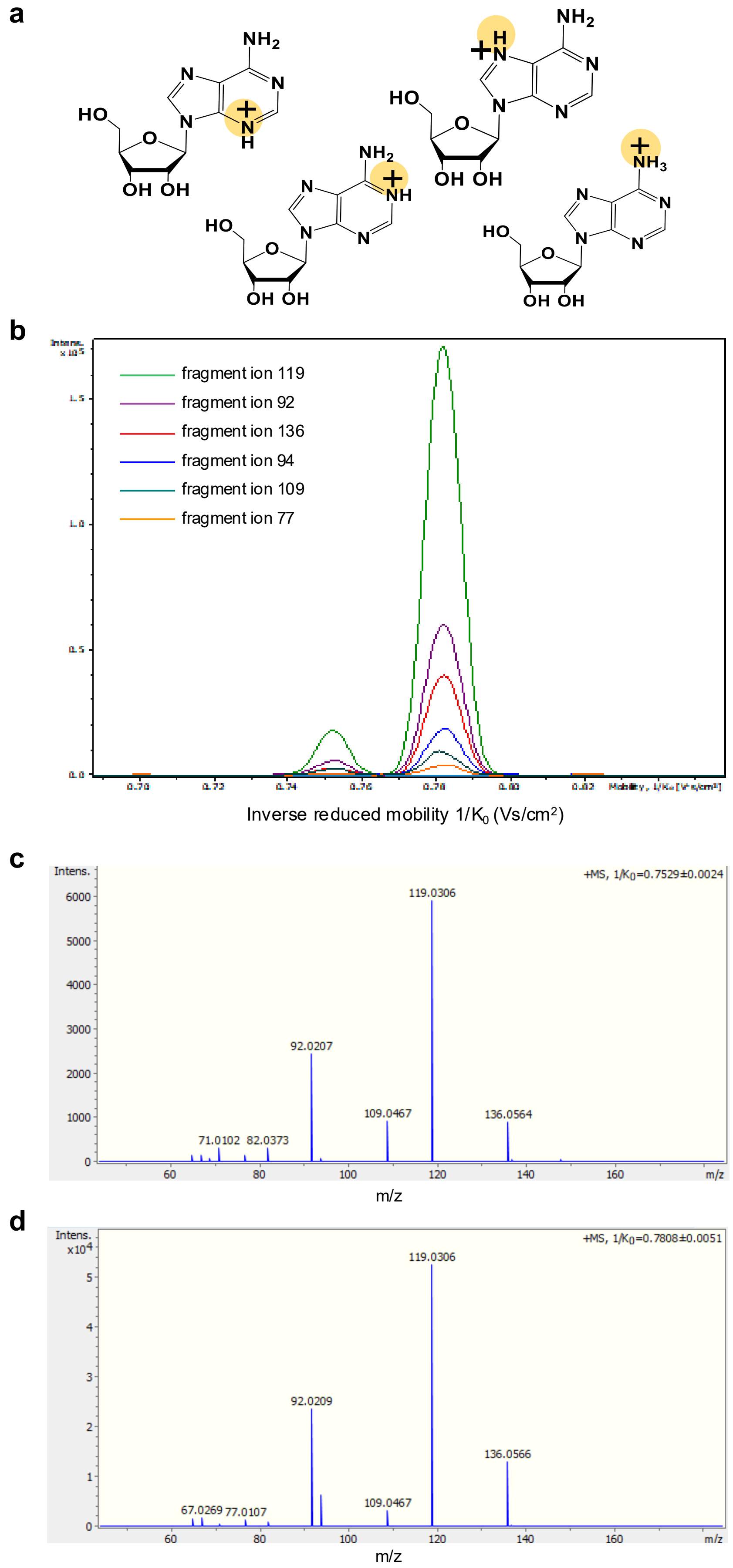


**Supplementary Figure 5. Charge isomers of adenosine resolved by TIMS.** a) Potential protonation sites of adenosine in positive mode. Two entries for the adenosine $[M+H]^+$ ion can be found in the Unified CCS Compendium with CCS values of 155.5 and 161.6, but no further reference data are available to differentiate charge isomers of adenosine. MS/MS fragmentation of the adenosine $[M+H]^+$ precursor ion posterior to TIMS separation shows that the two 4D peaks lead to different fragmentation patterns. b) EIMs of the fragment ions. c) MS/MS spectrum at the apex of the IMS peak at $1/K_0$ = 0.753. d) MS/MS spectrum at the apex of the IMS peak at $1/K_0$ = 0.781. Both MS/MS spectra share dominant fragment ions at *m/z* 92, 109, 119, and 136. However, only one MS/MS spectrum contains fragment ions at *m/z* 71, 82, and 94. Additionally, different ratios between the product ions are observed. We note that reference MS/MS data from public spectral libraries were not acquired by using IMS and therefore represent fragments from a distribution of charge isomers.

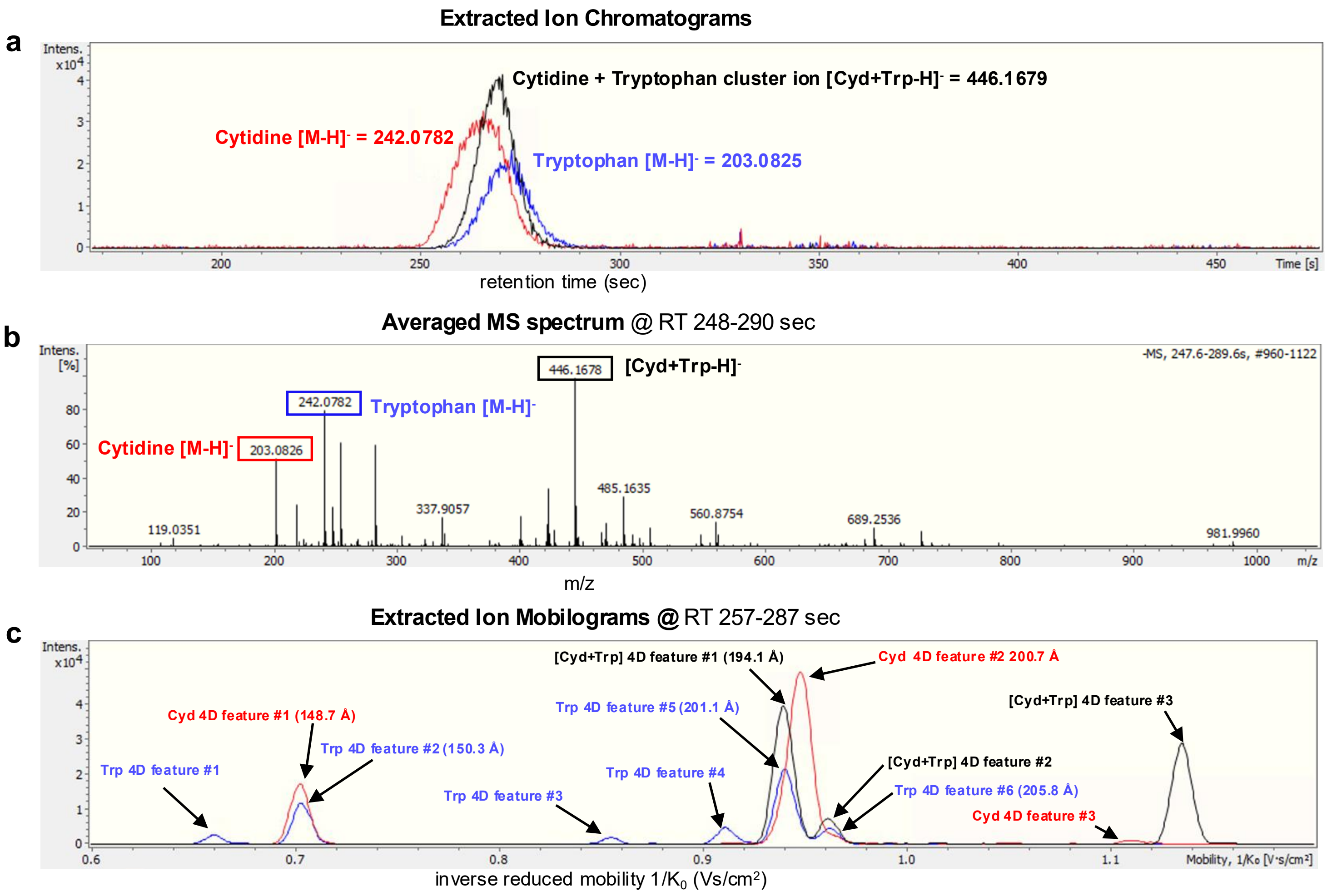


**Supplementary Figure 6**: **Confirmation of non-injective 4D features arising from post-IMS dissociation of ion clusters**. **a)** EICs of cytidine [M-H]-, tryptophan [M-H]-, and a cytidine-tryptophan ion cluster [Cyd+Trp-H]-. **b)** Averaged MS spectrum between retention time 257 and 287 seconds (scan #996 to scan #1114). **c)** EIMs of cytidine, tryptophan, and the cytidine-tryptophan cluster [M-H]- ions. The calculated CCS values of tryptophan 4D feature #2 (150.3 Å) and cytidine 4D feature #1 (148.7 Å) match the corresponding Unified CCS Compendium values of150.6 Å for tryptophan and 150 Å for cytidine (5% tolerance). Interestingly, we observed that 4D feature #1 and #2 of the [Cyd+Trp-H]- ion cluster co-migrated with tryptophan 4D features #5 and #6. The calculated CCS values of tryptophan 4D features #5 and #6 are 201.1 Å and 205.8 Å, which is more than a 30% difference than the library CCS value for Tryptophan. The co-migration pattern suggests a dissociation of the [Cyd+Trp-H]- ion cluster to produce a tryptophan ion with a neutral loss of a cytidine molecule.

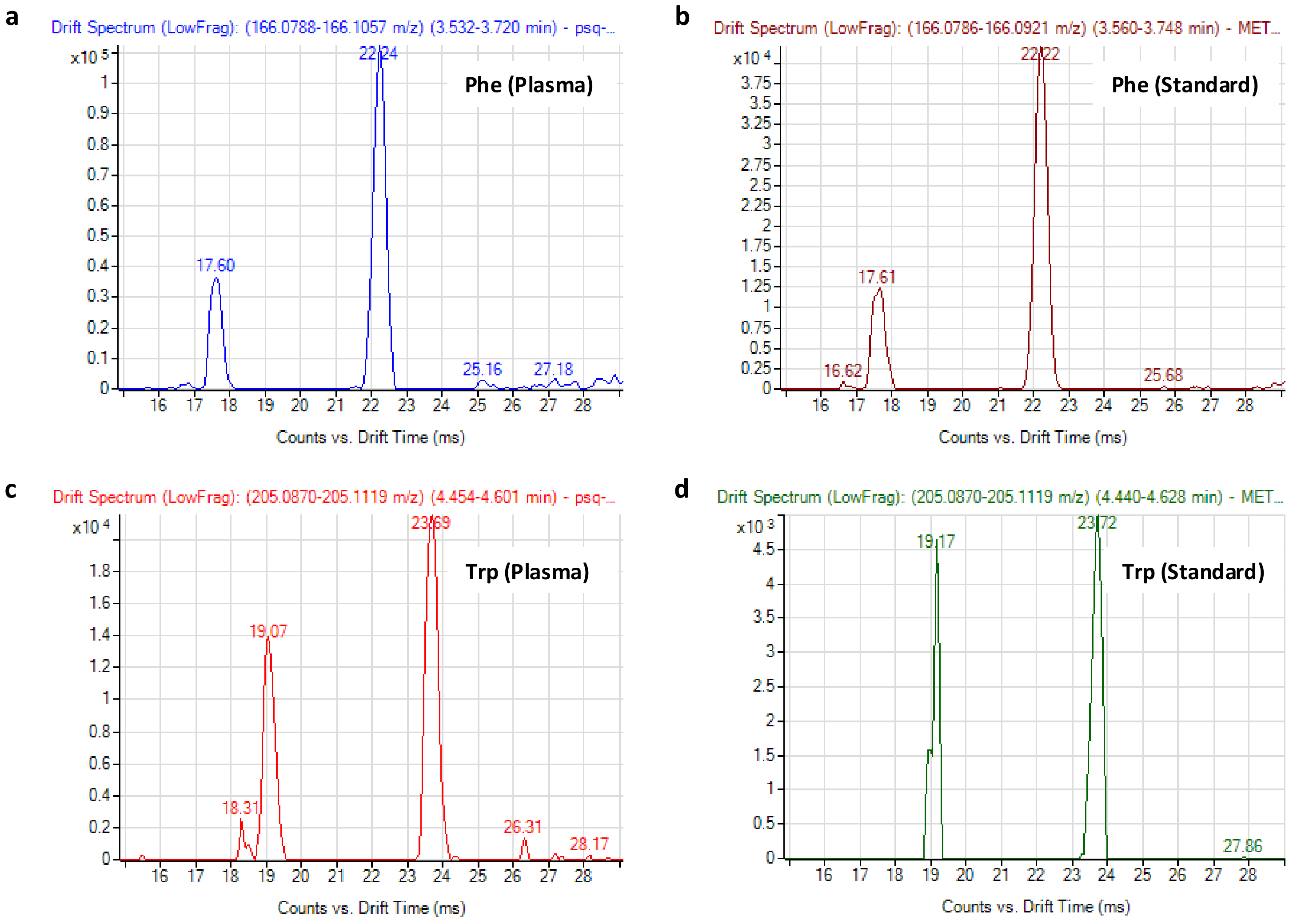


**Supplementary Figure 7**: **Non-injective 4D features for phenylalanine (Phe) and Tryptophan (Trp).** a) Phenylalanine has two non-injective 4D features in human plasma. b) A phenylalanine standard also produced two non-injective 4D features. c) Tryptophan has two non-injective 4D features in human plasma. d) A tryptophan standard also produced two non-injective 4D features. All data were collected with identical methods, using DTIMS-MS in positive mode. The ratios of the non-injective features measured from plasma are similar to those measured from each authentic chemical standard.

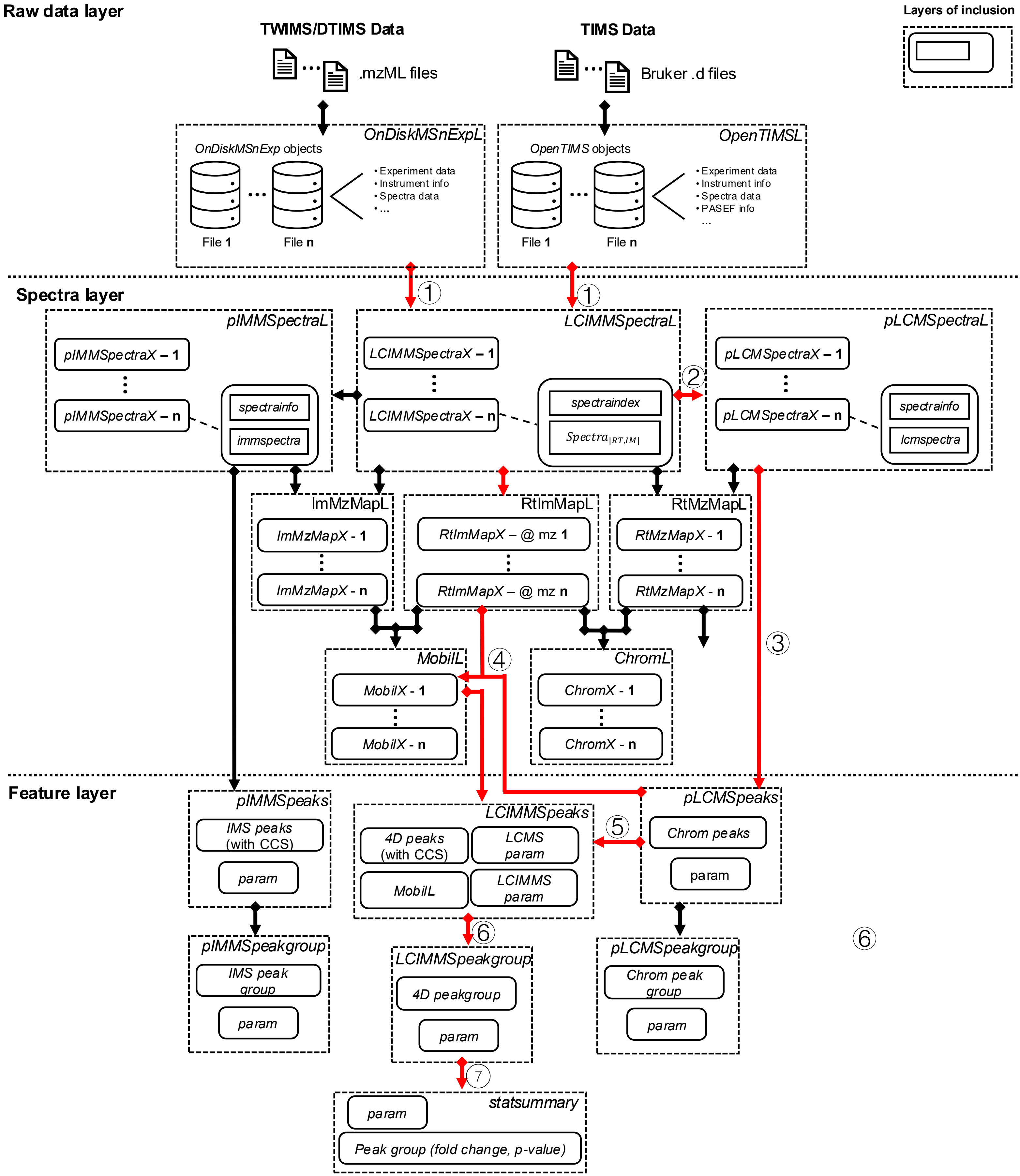


**Supplementary Figure 8: Data-processing workflow of XCiMS.** The processing architecture of XCiMS consists of a raw data layer, a spectra layer, and a feature layer. At the raw data layer, XCiMS applies the *MSnbase* R package to parse mzML files and the *opentimsr* R package to parse Bruker .d files into vendor-agnostic 4D spectra (*LCIMMSpectraL*). At the spectra layer, data objects suffixed with "*X*" are for single-file processing, while data objects suffixed with "*L*" contain multiple "*X*" type objects for multi-file processing. A typical processing workflow is highlighted in red with 6 steps detailed in Methods. Optionally, 3D intensity maps (RT-IM, IM-MZ, and RT-MZ) and 2D extracted ion traces (EIC, EIM) can be extracted at the spectra layer. At the feature layer, the *pLCMSpeaks* or *pIMMSpeaks* objects are constructed by detecting chromatographic and ion-mobility peaks from the *pLCMSpectraL or pIMMSpectraL* object. *LCIMMSpeaks* objects *for* 4D features are constructed by detecting IMS peaks of EIMs extracted from RT-IM ROIs. *pLCMSpeakgroup, pIMMSpeakgroup,* and LCIMM*Speakgroup* objects are optionally constructed by grouping *pLCMSpeaks, pIMMSpeaks,* or *LCIMMSpeaks* when sample class information is available.

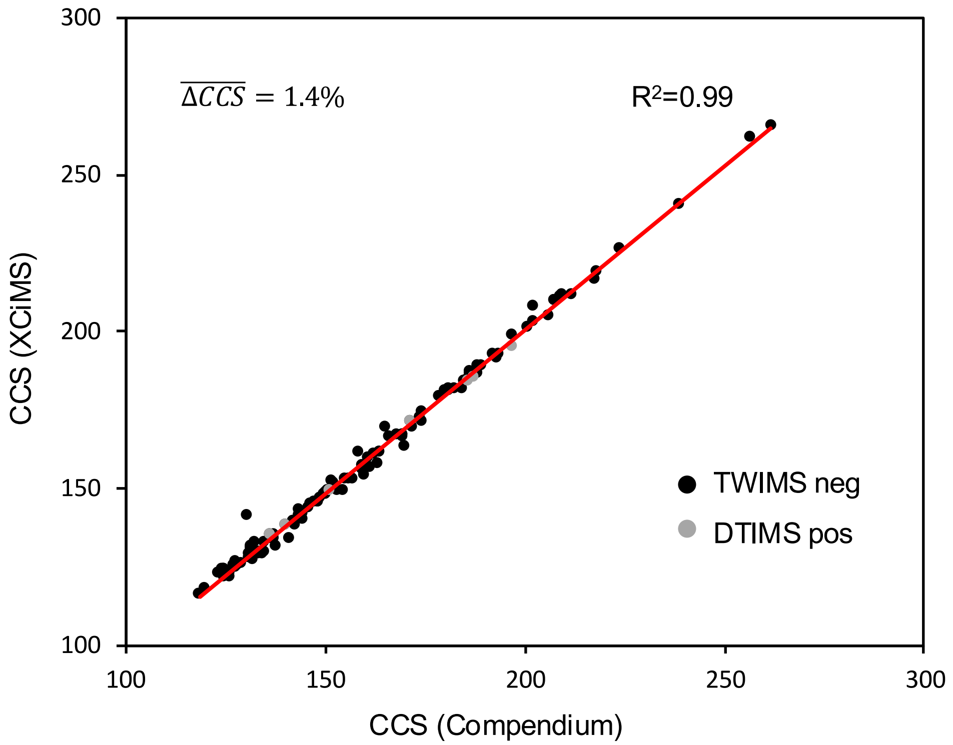


**Supplementary Figure 9**: **Calculating CCS for metabolite annotation**. Correlation between XCiMS-calculated CCS values and CCS values in the Unified CCS Compendium for polar metabolite standards (MSMLS). Phosphoenolpyruvate was excluded due to the wrong sum formula in the CCS Compendium (12/28/2024, index 2096). Coenzyme A is not shown. It was calculated as CCS = 193 in XCiMS and is 240 in the CCS Compendium.

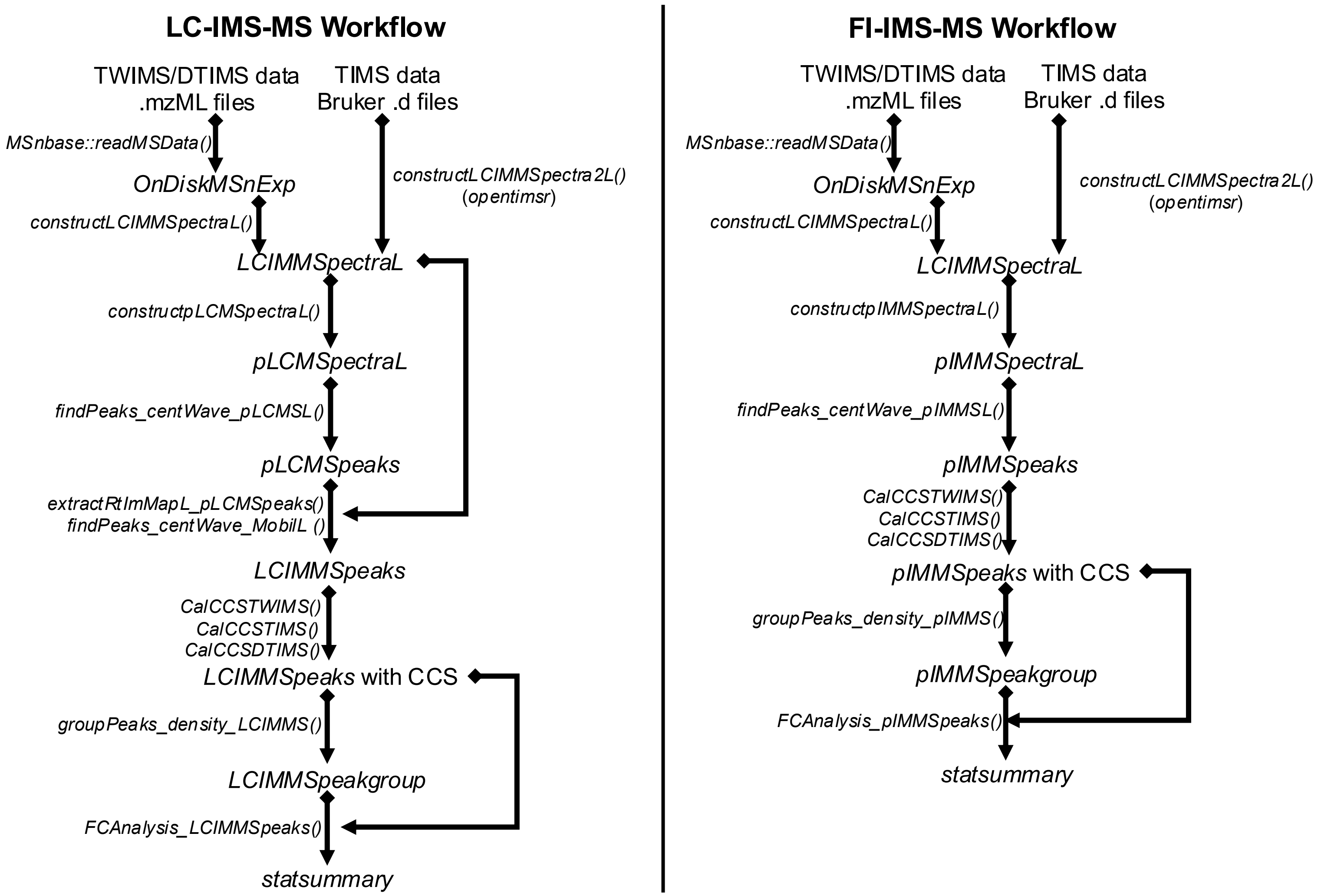


**Supplementary Figure 10:** XCiMS data-processing workflow for LC-IMS-MS and FI-IMS-MS experiments.
